# Neuronal morphologies built for reliable physiology in a rhythmic motor circuit

**DOI:** 10.1101/319582

**Authors:** Adriane G. Otopalik, Eve Marder

## Abstract

The neurons of the crustacean stomatogastric ganglion (STG) exhibit highly-conserved firing patterns, voltage waveforms, and circuit functions despite quantifiable animal-to-animal variability in their neuronal morphologies. In recent work, we showed that one neuron type, the Gastric Mill (GM) neuron, is electrotonically compact and operates much like a single compartment, despite having thousands of branch points and a total cable length on the order of 10 mm. Here, we explore how STG neurite morphology shapes voltage signal propagation and summation in four STG neuron types. We use focal glutamate photo-uncaging in tandem with somatic intracellular recordings to examine passive electrotonic structure and voltage signal summation in the GM neuron and three additional STG neuron types: Lateral Pyloric (LP), Ventricular Dilator (VD), and Pyloric Dilator (PD) neurons. In each neuron, we measured the amplitudes and apparent reversal potentials (E_rev_s) of inhibitory responses evoked with focal glutamate photo-uncaging at more than 20 sites varying in their distance (100–800 μm) from the somatic recording site in the presence of TTX. Apparent E_rev_s were relatively invariant (mean CVs = 0.04, 0.06, 0.05, and 0.08 for 5–6 GM, LP, PD, VD neurons, respectively), suggesting that all four neuron types are similarly electrotonically uniform and compact. We then characterized the directional sensitivity and arithmetic of voltage summation (with fast sequential activation of 4–6 sites) in individual STG neurites. All four neuron types showed no directional bias in voltage signal summation and linear voltage summation. We motivate these experiments with a proof-of-concept computational model that suggests the immense tapering of STG neurite diameters: from 10–20 μm to sub-micron diameters at the terminal tips, may explain the uniform electrotonic structures experimentally observed and contribute to the robust nature of this central pattern-generating circuit.

## Introduction

Neurons often present complex and highly-branched morphologies. The distributed biophysical cable properties arising from neuronal geometry influence current flow and voltage signal propagation; like a leaky cable, ease of voltage signal propagation is a direct consequence of neurite diameter and membrane resistance, and is inversely related to axial resistance (Rall, 1959, 1960, 1969; Jack et al., 1975). How these basic biophysical principles play out in a functioning neuron is dependent upon where synaptic inputs, receptors, and ion channels, which may shunt or amplify this signal, are distributed across the path of propagation (London and Häusser, 2005). Of course, these properties vary across neuron types. Therefore, it is critical to characterize the passive physiology of diverse neuron types to uncover the breadth of biophysical organizations utilized in diverse neuron types and circuit contexts.

In two recent studies, we characterized the morphology (Otopalik et al., 2017a) and passive physiology (Otopalik et al., 2017b) of the identified neurons of the crustacean stomatogastric ganglion (STG), a small central pattern-generating circuit mediating the rhythmic contractions of the animal’s foregut. The 14 identified neurons of the STG present distinct, cell-type-specific physiological waveforms, firing patterns and circuit functions. We quantified numerous morphological features pertaining to the macroscopic branching patterns and fine cable properties of four neuron types (Otopalik et al., 2017a). There was quantifiable inter-animal variability in all features within neuron types, and no single metric or combination of metrics distinguished the four neuron types. We then asked: How do STG neurons produce reliable firing patterns, given their variable morphologies across animals? As a first examination of how morphology maps to physiology in the STG, we characterized the electrotonic structure of one STG neuron type, the Gastric Mill (GM) neuron. We found that GM neurons are surprisingly electrotonically compact, despite their expansive and complex neuronal structures (Otopalik et al., 2017b). We suggested that compact electrotonic structures may effectively mask the physiological consequences of morphological variability observed in GM neurons across animals (Otopalik et al., 2017a, b).

In the present study, we complement a proof-of-concept computational model with glutamate photo-uncaging experiments in single STG neurons to explore how STG neuronal morphology may yield compact electrotonic structures and uniform voltage signal propagation across the neurite tree.

## Results

### Equalizing Neurite Morphologies

In previous work, we were surprised to find that STG neurons exhibit neurites as long as 1 mm in length, and neurite diameters that often tapered from 10–20 microns at the primary neurite junction, to sub-micron diameters at the terminating tips (Otopalik et al., 2017a). Passive cable theory, first applied to the study of neurons by Wilfred Rall and colleagues, suggests that voltage signals passively propagating such long distances are likely to undergo a great deal of attenuation (Rall, 1960, 1964, 1969; Jack et al., 1975). This distance-dependent attenuation of voltage events leads to asymmetrical responses at the soma when multiple sites on the same dendrite, or neurite, are activated in fast sequence in the inward or outward directions (Rall, 1964; Gulledge et al., 2005). The simulation shown in Figure 1A-C recapitulates this phenomenon in a passive cable model reminiscent of that used by Rall (1964; Figure 1A). Inhibitory voltage events (Figure 1B, top) were evoked at five sites and activated in sequence in the inward (red) or outward (blue) directions (Figure 1B, bottom). Activation in the inward direction resulted in modest increases in response integral and peak (increase of 0.1 mV*s and 0.4 mV, respectively). This directional bias (inward integral minus the outward integral) and the arithmetic of voltage signal summation (inward response integral minus the linear sum of individual response integrals shown in Figure 1B, top) was dependent on the passive biophysical properties of the neurite (Figure 1C). As the membrane resistivity (R_m_) decreases, and the cable becomes increasingly leaky, this inward bias becomes less apparent and the voltage summation transitions from sublinear to linear. Likewise, increasing axial resistivity (R_a_) results in less directional bias and a transition from sublinear to linear summation.

**Figure 1.**
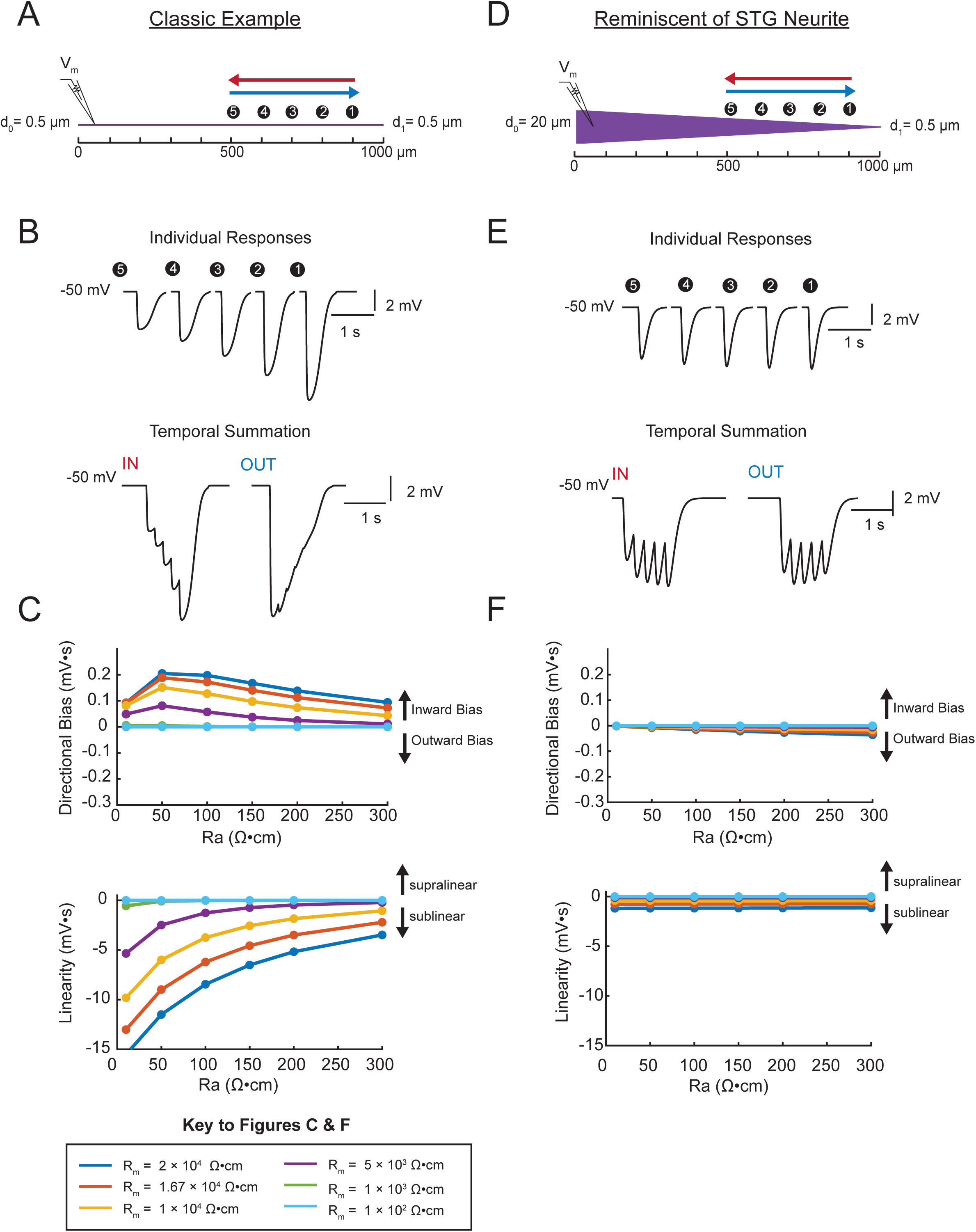
Simulating voltage summation in neurites with diverse passive properties and geometries. Voltage summation experiments were simulated in NEURON. Inhibitory potentials were evoked at 5 sites (numbered) individually or in sequence at 5 Hz in the inward or outward directions relative to the recording site. A-C summarizes results from a classic cable model without a tapering geometry. D-F summarizes results from a cable model with neurite geometry reminiscent of that observed in STG neurons. **A**. Schematic showing a classic cable model simulation for a 0.5 μm-diameter cable. **B**. Simulated traces for activation of individual sites (top) and sequential activation in either direction (below), as depicted by colored arrows in A. **C**. Quantification of directional bias and linearity for 0.5 μm-diameter cables varying in their passive properties. Top: directional bias as a function of specific axial resistivity (R_a_ in Ω*cm). Directional bias was calculated as inward integral minus the outward integral; positive values suggest an inward bias, whereas negative values suggest an outward bias. Points close to the y = 0 suggest no directional preference. Bottom: Linearity as a function of R_a_. Linearity was calculated as the inward integral minus the integral of the arithmetic sum of events evoked at individual sites (as in B, top); positive values suggest supralinear summation, whereas negative values suggest sublinear voltage summation. Points close to the y = 0 suggest linear summation. All plots show data for a range of specific membrane resistivity (R_m_ in Ω*cm^2^) values in different colors (indicated in the key below). Data for the full simulation exploring a broad parameterspace are shown in Supplements 1 and 2 to Figure 6. **D**. Schematic showing a cable model simulation for a neurite that tapers from 20 μm at the recording end (d_0_) to 0.5 μm at the distal end (d_1_). E. Simulated traces for activation of individual sites (top) and sequential activation in either direction (below), as depicted by colored arrows in D. **F**. Directional bias and linearity plotted as a function of R_a_ and varying R_m_ values (indicated in key) for a cable with the geometry shown in D. B and E: R_m_ = 10000 Ω*cm^2^ and R_a_ = 100 Ω*cm.

To explore how neurite geometry may also shape voltage summation, we executed this simulation in a modest library of 720 cable models with varying geometries and passive properties (R_a_ and R_m_ values; see Materials and Methods). Interestingly, we found that directional bias was abolished and that voltage summation was consistently linear across a broad range of R_a_ and R_m_ values in neurites with geometries reminiscent of those observed in STG neurites (Figures 1D-F). Directional bias and linearity for the full model library can be found in Supplements 1 and 2 to Figure 1. Taken together, this simulation suggests that the immense tapering of neurites likely contributes to the uniform electrotonic structures observed in Gastric Mill (GM) neurons, one STG neuron type (Otopalik et al., 2017b).

In the following experiments, we test this hypothesis and the generalizability of this morphological solution in four STG neuron types using focal glutamate photo-uncaging in tandem with electrophysiological recordings at the soma. First, we will probe the electrotonic structures of four STG neuron types by measurement of apparent reversal potentials (E_rev_s) for inhibitory events evoked across the neurite tree (as in Otopalik et al., 2017b). Then, we will assess voltage propagation and summation arithmetic by evoking sequential inhibitory events on single neurites, thereby mimicking the same stimuli depicted in our computational simulation.

### Compact Electrotonic Structures in Four STG Neuron Types

Here, we broadly characterize the electrotonic structure of three additional STG neuron types: Lateral Pyloric (LP), Ventricular Dilator (VD), and Pyloric Dilator (PD), as well as corroborate our previous findings in the GM neuron. These four neuron types present equally complex and variable morphologies, but distinct voltage waveforms and circuit functions. The PD, LP, VD, and GM neurons were unambiguously identified by their innervation patterns (Figure 2A) and by matching their intracellular spiking patterns with concurrent extracellular recordings of nerves known to contain their axons (Figure 2B). PD and LP neurons innervate two muscles in the pylorus of the foregut (Figure 2A) and participate in the ongoing, triphasic pyloric rhythm (Figure 2B). PD and LP can both be identified by matching their intracellular firing patterns with spiking units on the lateral ventricular nerve (*lvn*). The VD neuron innervates the cv1 muscle of the pylorus and can be identified on the medial ventricular nerve (*mvn*). The GM neuron participates in the episodic gastric mill rhythm, innervates gm1, 2, and 3 muscles, and can be identified on the dorsal gastric nerve (*dgn*). When filled with fluorescent dye, each neuron presents highly branched and expansive neurite trees (Figure 2C). The morphological features of these neuron types have been described quantitatively and in detail in previous studies (Wilensky et al. 2003; Bucher et al., 2007; Thuma et al. 2009; Otopalik et al., 2017a).

**Figure 2.**
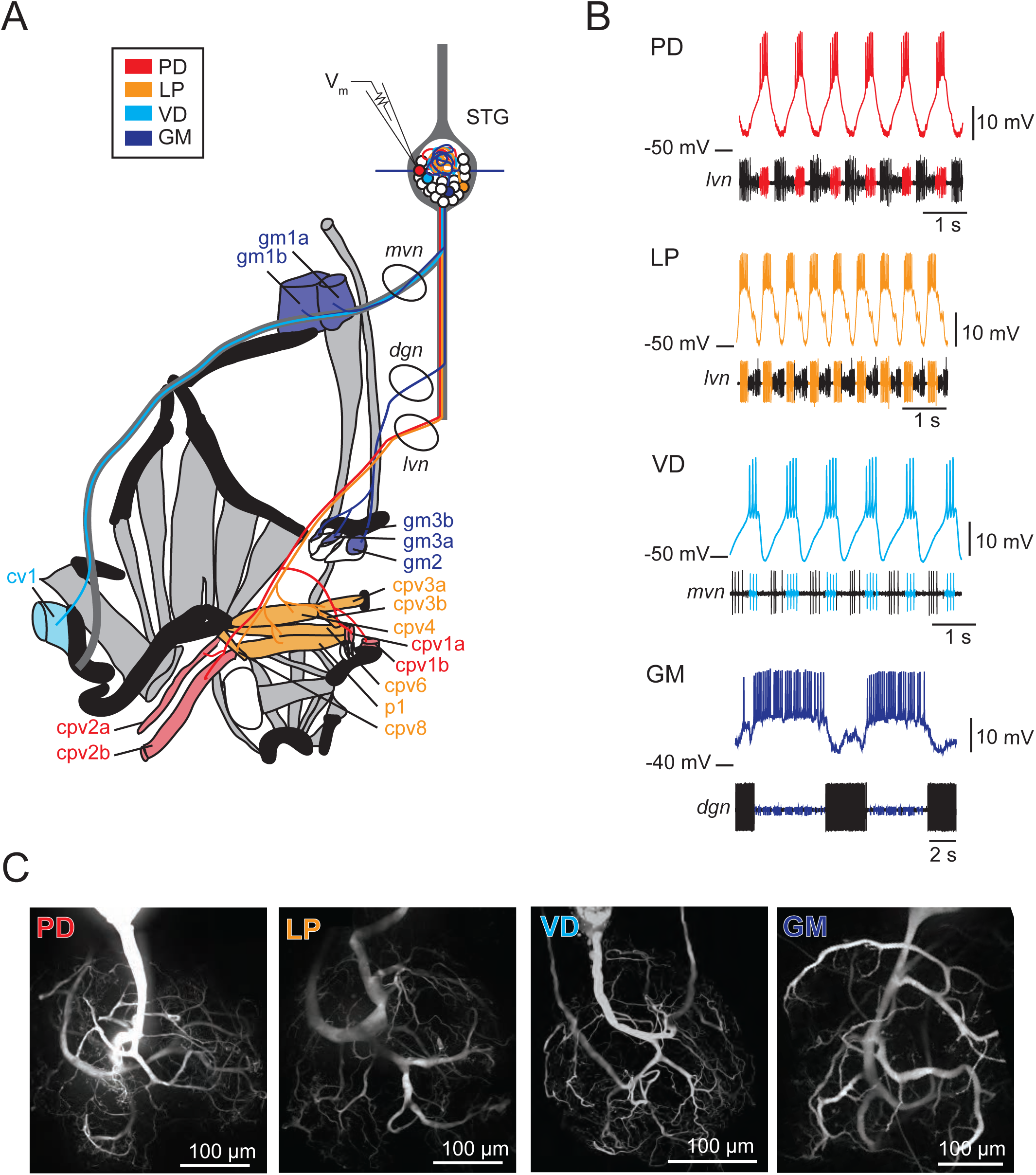
Characteristics of four identified STG neuron types. A. Bilateral innervation patterns of each neuron type, depicted on one side of the dissected foregut. Axons from the four neuron types project from the STG (top) and project to specific muscle groups (indicated with the same colors). The axonal spiking activity for each of these neurons can be recorded with extracellularly nerve recordings at the circled locations on the *mvn, dgn*, and *Ivn* nerves. B. Each neuron type can be identified physiologically by matching intracellular firing patterns with concurrent extracellular recordings of nerves known to contain their axons (as shown in A). C. Representative z-projections show complex neurite trees for each neuron type acquired at 40x magnification. PD = Pyloric Dilator; LP = Lateral Pyloric; VD = Ventricular Dilator; GM = Gastric Mill; *Ivn* = lateral ventricular nerve; *dgn* = dorsal gastric nerve; *mvn* = medial ventricular nerve. For all subfigures: red = PD, orange = LP; light blue = VD; dark blue = GM.

We probed passive voltage signal propagation by evoking inhibitory potentials at numerous sites on the neurite tree with focal photo-uncaging of MNI-glutamate and two-electrode current clamp recordings at the soma (as in Otopalik et al., 2017b). Figure 3A shows example traces of evoked inhibitory potentials at six sites on an individual branch of a PD neuron. When the somatic membrane potential is at rest (approximately −50 mV) evoked events at the six sites are inhibitory, but vary in their magnitude. Two-electrode current clamp was used to manipulate the somatic membrane potential (between −40 and −100 mV) and the apparent E_rev_s for events at each site were determined by plotting response amplitude as a function of somatic membrane potential (Figure 3B). The x-intercept of the linear fit of these data serves as our measure of the apparent E_rev_ for each site. Across the six sites evaluated in Figure 3A and B, apparent E_rev_s ranged between −59 and −70 mV.

**Figure 3.**
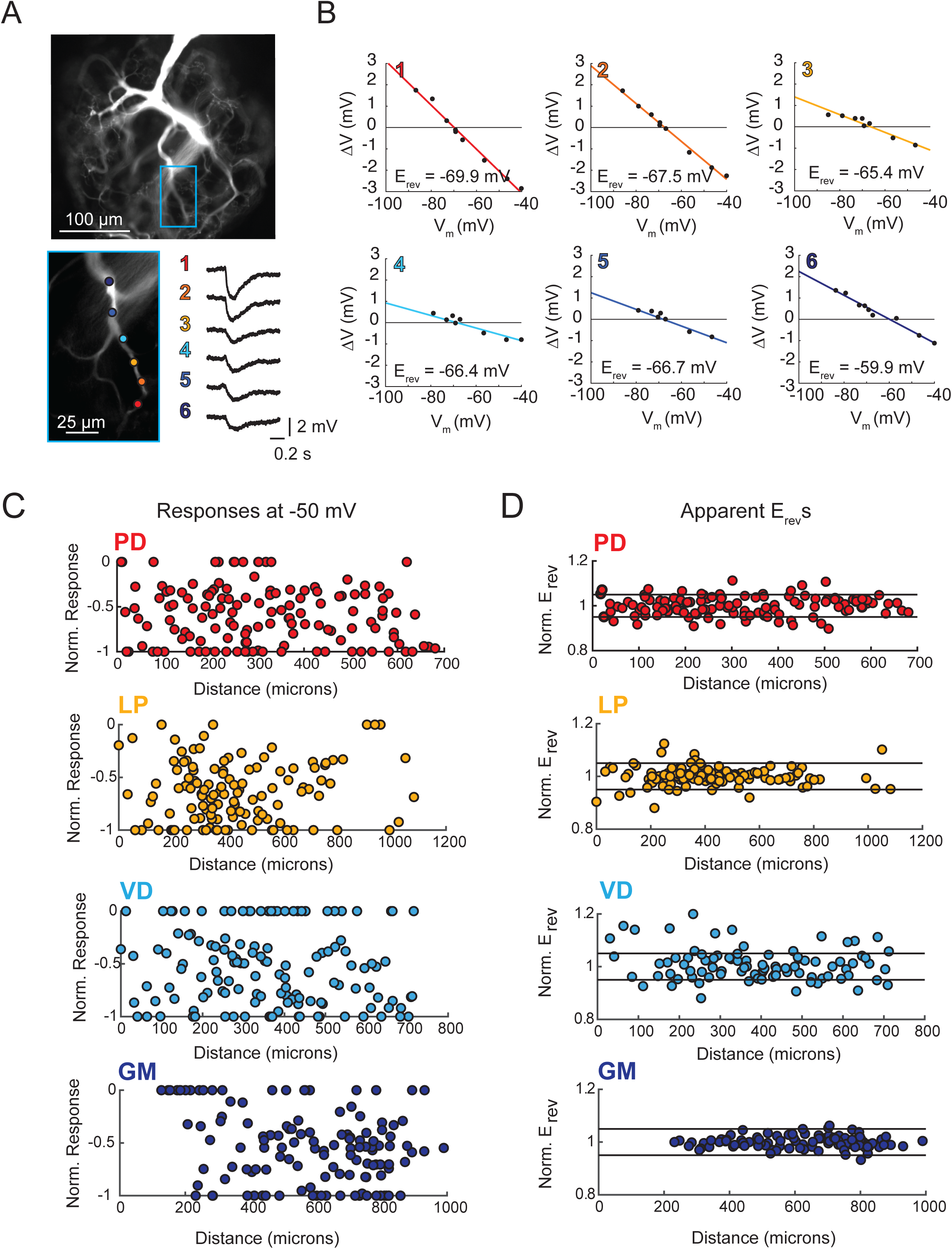
Variable response amplitudes and invariant apparent reversal potentials (E_rev_s) across STG neuronal structures. A and B: Glutamate photo-uncaging and measurement of apparent E_rev_s in a representative PD neurite. A. Fluorescence images of an Alexa Fluor 488 dye-fill showing the neurite tree at 20x magnification and a single branch at 40x magnification (outlined in blue on the 20x image). Glutamate photo-uncaging sites are indicated with colored circles and corresponding responses (measured at the soma) for each site are shown. At each site, inhibitory events were evoked with a 1-ms, 405-nm laser pulse at a starting somatic membrane potential of −50 mV. B. Plots showing evoked response amplitude (∆V) as a function of somatic membrane potential (V_m_). At each site, glutamate responses were evoked at varying somatic membrane potentials (achieved with two-electrode current clamp). These data were fit with a linear regression (lines) and the E_rev_ for each site was calculated as the x-intercept of this fit (values for each site shown on bottom right of each plot). C. Response amplitudes plotted as a function of distance from the somatic recording site for each neuron type. Maximum response amplitudes were measured at −50 mV for individual sites and normalized to the maximum response amplitude within each neuron (−1 is equivalent to the maximum response within individual neurons). There was no quantitative relationship between response amplitude and distance (supported by poorly fit linear regression analyses in Table 1). D. Apparent E_rev_s for each site were normalized to the mean apparent E_rev_s within each neuron (1 is representative to the mean). Horizontal black lines denote boundaries of + 5% of the mean apparent E_rev_s and serve as a graphical depiction of the low variance in apparent E_rev_s within each neuron for sites as far as 800–1000 μm away from the soma. Raw response amplitudes and apparent E_rev_s for individual sites in individual neurons are shown in Supplements 1–4 to Figure 3.

Maximal response amplitudes (as measured at a somatic membrane potential of −50 mV) and apparent E_rev_s were measured for numerous sites across the neurite trees of each neuron type (for 10–30 distinct sites across the neurite trees of five LP, VD, GM neurons and six PD neurons; Tables 1 and 2). These data are summarized in Figures 3C and D, where maximal response amplitudes and apparent E_rev_s for individual sites are plotted as a function of distance from the somatic recording site for each neuron. For comparison across multiple neurons, maximal response magnitudes (which varied between 0–4 mV) were normalized to the minimum (always 0 mV) and maximum response amplitudes within each neuron (Supplements 1–4 to Figure 3 show raw maximum response amplitudes and apparent E_rev_s for individual PD, LP, VD, and GM neurons, respectively). Figure 3C shows that, across all neuron types, maximal response amplitudes show no quantitative trend as a function of distance, nor is there any evidence of normalization of response amplitude with distance (such that response amplitudes are uniform across sites). This is reflected in poorly-fit and insignificant linear regressions of these data (Table 1).

**Table 1.** Linear regression analyses for response amplitudes and apparent reversal potentials (E_rev_s) as a function of distance from the somatic recording site for sites in individual neurons or pooled by cell type. The data contributing to these analyses are shown graphically in Figure 3C and 3D and Supplements 1–4 to Figure 3.

| Neuron | Sites | Amplitude vs. Distance | | | | Apparent $E_{rev}$ vs. Distance | | | |
| --- | --- | --- | --- | --- | --- | --- | --- | --- | --- |
| | | MSE | R | p | slope (mV/μM) | MSE | R | p | slope (mV/μM) |
| PD | 30 | 0.08 | 0.05 | 7.98E-01 | 1.67E-04 | 13.61 | 0.17 | 4.22E-01 | -7.23E-03 |
| PD | 23 | 0.15 | -0.05 | 8.38E-01 | -1.03E-04 | 4.66 | -0.73 | 8.43E-05 | -1.35E-02 |
| PD | 10 | 0.05 | -0.06 | 8.28E-01 | -1.77E-04 | 6.65 | -0.42 | 2.26E-01 | -1.60E-02 |
| PD | 23 | 0.37 | -0.47 | 2.05E-02 | -1.56E-03 | 15.13 | -0.19 | 3.88E-01 | -3.87E-03 |
| PD | 20 | 0.40 | -0.18 | 4.26E-01 | -6.35E-04 | 17.96 | 0.05 | 8.50E-01 | 1.05E-03 |
| PD | 23 | 0.77 | 0.40 | 5.18E-02 | 2.62E-03 | 32.09 | 0.37 | 8.55E-02 | 1.62E-02 |
| **PD<sub>MEAN</sub>** | **21.5** | **0.30** | **-0.05** | **4.94E-01** | **5.18E-05** | **11.60** | **-0.29** | **3.77E-01** | **-7.91E-03** |
| **PD<sub>SD</sub>** | **6.5** | **0.27** | **0.29** | **3.86E-01** | **1.40E-03** | **9.79** | **0.38** | **3.04E-01** | **1.16E-02** |
| LP | 11 | 0.14 | 0.43 | 1.23E-01 | 5.87E-04 | 12.09 | -0.81 | 2.67E-03 | -1.54E-02 |
| LP | 26 | 0.50 | -0.02 | 9.25E-01 | -1.15E-04 | 5.07 | -0.19 | 3.08E-01 | -4.00E-03 |
| LP | 25 | 0.17 | -0.15 | 4.75E-01 | -5.56E-04 | 14.18 | -0.17 | 4.16E-01 | -5.79E-03 |
| LP | 25 | 0.34 | -0.40 | 4.70E-02 | -1.56E-03 | 5.75 | -0.09 | 6.75E-01 | -2.15E-03 |
| LP | 24 | 0.51 | 0.02 | 9.30E-01 | 7.01E-05 | 0.70 | 0.55 | 5.00E-03 | 7.26E-03 |
| **LP<sub>MEAN</sub>** | **22.2** | **0.33** | **-0.02** | **5.00E-01** | **-3.15E-04** | **10.16** | **-0.14** | **2.81E-01** | **-4.02E-03** |
| **LP<sub>SD</sub>** | **6.3** | **0.18** | **0.30** | **4.22E-01** | **8.08E-04** | **5.45** | **0.48** | **2.86E-01** | **8.12E-03** |
| VD | 27 | 0.31 | 0.05 | 7.95E-01 | 2.32E-04 | 193.71 | -0.35 | 7.64E-02 | -4.17E-02 |
| VD | 19 | 0.54 | -0.53 | 3.49E-03 | -6.51E-03 | 28.63 | -0.45 | 5.57E-02 | -3.85E-02 |
| VD | 24 | 0.28 | -0.14 | 5.06E-01 | -2.46E-04 | 41.44 | 0.22 | 3.16E-01 | 5.07E-03 |
| VD | 15 | 0.07 | -0.20 | 3.36E-01 | -3.28E-04 | 15.74 | -0.29 | 2.97E-01 | -7.76E-03 |
| VD | 15 | 0.87 | 0.35 | 4.73E-02 | 1.81E-03 | 33.58 | -0.69 | 3.29E-03 | -5.59E-02 |
| **VD<sub>MEAN</sub>** | **20.0** | **0.42** | **-0.09** | **3.38E-01** | **-1.01E-03** | **62.62** | **-0.31** | **1.50E-01** | **-2.78E-02** |
| **VD<sub>SD</sub>** | **5.4** | **0.31** | **0.33** | **3.29E-01** | **3.19E-03** | **73.87** | **0.33** | **1.46E-01** | **2.54E-02** |
| GM | 10 | 0.04 | -0.84 | 6.83E-04 | -3.23E-03 | 0.56 | -0.14 | 6.93E-01 | -1.17E-03 |
| GM | 28 | 0.09 | -0.53 | 2.02E-03 | -8.74E-04 | 2.09 | 0.31 | 1.13E-01 | 2.49E-03 |
| GM | 25 | 0.64 | -0.24 | 1.78E-01 | -1.12E-03 | 5.56 | -0.13 | 5.28E-01 | -2.23E-03 |
| GM | 20 | 0.09 | -0.41 | 3.50E-02 | -5.76E-04 | 11.76 | 0.17 | 4.76E-01 | 9.37E-03 |
| GM | 19 | 0.03 | -0.57 | 4.56E-03 | -1.14E-03 | 4.03 | -0.24 | 3.26E-01 | -6.53E-03 |
| **GM<sub>MEAN</sub>** | **20.4** | **0.18** | **-0.52** | **4.41E-02** | **-1.39E-03** | **4.80** | **-0.01** | **4.27E-01** | **3.85E-04** |
| **GM<sub>SD</sub>** | **6.9** | **0.26** | **0.22** | **7.62E-02** | **1.06E-03** | **4.33** | **0.23** | **2.19E-01** | **5.97E-03** |

**Table 2.** Apparent Mean E_rev_s, standard deviations (SD) and coefficients of variance (CV) for individual neurons and within neuron type. Columns 2 and 3 indicate the number of individual sites evaluated and the number of separate branches probed within each neuron.

| Neuron | Sites | Branches | Mean $E_{rev}$<br>(mV) | SD<br>(mV) | CV |
| --- | --- | --- | --- | --- | --- |
| PD | 30 | 6 | -64.70 | 3.80 | 0.06 |
| PD | 23 | 4 | -66.30 | 3.20 | 0.05 |
| PD | 10 | 3 | -74.30 | 3.00 | 0.04 |
| PD | 23 | 4 | -64.18 | 4.05 | 0.06 |
| PD | 20 | 4 | -69.56 | 4.35 | 0.06 |
| PD | 23 | 4 | -83.00 | 6.22 | 0.08 |
| **PD<sub>MEAN</sub>** | **21.5** | **4.2** | **-70.34** | **4.10** | **0.06** |
| LP | 11 | 2 | -65.72 | 6.18 | 0.09 |
| LP | 26 | 5 | -72.82 | 2.33 | 0.03 |
| LP | 25 | 5 | -71.87 | 3.89 | 0.05 |
| LP | 25 | 5 | -81.67 | 4.07 | 0.05 |
| LP | 24 | 5 | -71.87 | 3.89 | 0.05 |
| **LP<sub>MEAN</sub>** | **22.2** | **4.4** | **-72.79** | **4.07** | **0.05** |
| VD | 27 | 4 | -76.22 | 3.75 | 0.05 |
| VD | 19 | 4 | -80.83 | 6.14 | 0.08 |
| VD | 24 | 4 | -83.86 | 6.75 | 0.08 |
| VD | 15 | 4 | -74.81 | 4.29 | 0.06 |
| VD | 15 | 5 | -77.84 | 8.23 | 0.11 |
| **VD<sub>MEAN</sub>** | **20** | **4.2** | **-78.71** | **5.83** | **0.08** |
| GM | 10 | 2 | -62.33 | 0.80 | 0.01 |
| GM | 28 | 5 | -74.00 | 1.55 | 0.02 |
| GM | 25 | 6 | -74.78 | 2.43 | 0.03 |
| GM | 20 | 5 | -75.42 | 3.57 | 0.05 |
| GM | 19 | 4 | -65.66 | 7.05 | 0.11 |
| **GM<sub>MEAN</sub>** | **20.4** | **4.4** | **-70.44** | **3.08** | **0.04** |

Figure 3D shows apparent E_ev_s as a function of distance from the somatic recordings site. For comparison of responses evoked at sites on multiple neurons, apparent E_rev_s were normalized to and plotted as a percent of the mean E_rev_ across sites within each neuron. Horizontal lines denote 0.05, or 5%, above and below the mean E_rev_. For each neuron type, there appears to be no substantial hyperpolarization in apparent E_rev_s with distance from the somatic recording site (this is validated with linear regressions shown in Table 1). Figure 3B shows that GM neurons present exceptionally invariant apparent E_rev_s across sites on their neurite trees (mean coefficient of variance (CV) within individual GM neurons was 0.04; Table 2). This result is consistent with previous findings (Otopalik et al., 2017b). Likewise, LP, PD, and VD exhibit relatively invariant apparent Erevs (mean CVs were: 0.06, 0.05, and 0.08, respectively; Table 2). Although, it should be noted that a handful of VD neurons show higher standard deviations and CVs. Statistical comparison of CVs (ANOVA, [F(3, 17)=1.2 p=0.341]) and mean apparent E_rev_s (ANOVA, [F(3, 17)=2.29, p=0.1154]) revealed no statistically significant differences across neuron types. This suggests that the four neuron types are similarly electrotonically compact neuronal structures, wherein apparent E_rev_s typically varied by ≤ 10% of the mean within individual neurons. This translates to a range of mean E_rev_ + 6–8 mV within each individual neuron.

### Direction insensitivity of voltage signal propagation

Given the compact nature of the STG neurons assessed here and the immense tapering in neurite geometries previously observed (Otopalik et al., 2017a), we hypothesized that their neurites may be indifferent to direction of activation (described in Figure 1 and explored in Supplement 2 to Figure 1). To test for directional bias in voltage signal propagation, sequential voltage events were evoked at multiple sites within the same secondary branches (from tip to primary neurite junction; Figure 4Ai). The integrals of the summed responses for inward and outward activation were measured at the soma (Figure 4Aii). Directional preference for each branch was assessed by plotting the response integrals for the inward and outward directions against each other and comparison with the identity line, which is indicative of direction insensitivity wherein the inward and outward response integrals are equal (Figure 4Aiii). Any branches with points left of the identity line present an inward, or centripetal bias, whereas any points right of the identity line are suggestive of a centrifugal, or outward, bias. Interestingly, all four neuron types show little directional selectivity (this is supported by root-mean-square error values (RMSE) < 0.5 mV*s, a measure of goodness-of-fit to the identity line). Example traces and direction selectivity plots for each cell type can be found in Supplements 1–4 to Figure 4. Taken together, these results suggest that neurites in each of these four STG neuron types do not exhibit robust directional selectivity as has been observed in other cell types (London and Häusser, 2005).

**Figure 4.**
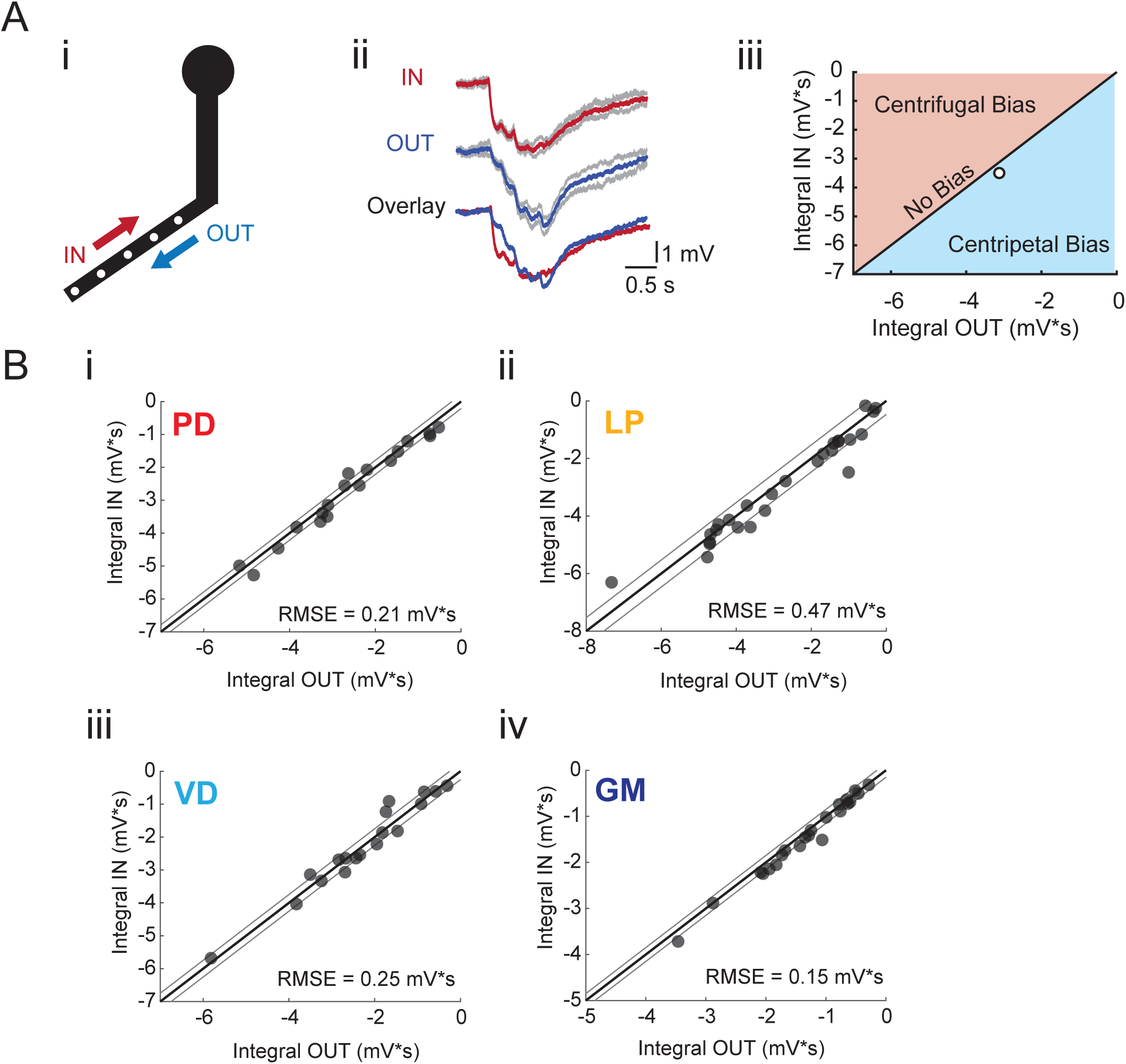
Directional sensitivity of voltage propagation in four neuron types. (i) 4–6 sites on single neurites were sequentially photo-activated at 5 Hz in the inward (IN) or outward (OUT) directions. The integrals (ii) of these inhibitory summation responses were calculated as the area above the trace (in mV*s) and plotted against each other as shown in (iii). As plotted, any points that lie to the right of the identity line (shaded in blue) show a centripetal, or inward bias, whereas any points that lie to the left of the identity line (shaded in red) show an outward, or centrifugal, bias. Any points near or on the identity line are unbiased (as is the case with the example traces shown in (ii), depicted with the white data point in (iii)). B. Directional bias plots for branches for numerous branches within neuron type: (i) 19 branches from 6 PD neurons, (ii) 27 branches from 9 LP neurons, (iii) 20 branches from 5 VD neurons, (iv) 22 branches from 5 neurons. These data were fit to the identity line, and the root-mean-square error (RMSE) boundaries for this fit is plotted in gray lines.

### Arithmetic of voltage signal integration

If voltage signal propagation in STG neurons is predominantly shaped by passive properties and tapered neurite geometries, and less so by active biophysical properties, which may shunt or amplify propagating voltage signals, we would expect to observe linear voltage summation as is predicted by our simulation (Figure 1 and Supplement 2 to Figure 1). To assess the arithmetic of voltage summation, we calculated the arithmetic sum of responses evoked at individual sites across single neurites (Figure 5Ai-iii; offset by 200 ms to mimic a 5-Hz sequential activation rate). The integrals of the measured response and expected arithmetic sum for a given branch were plotted against each other and compared with the identity line, which is indicative of linear summation, wherein the measured and expected response integrals are equal (Figure 5Aiv). Any branches with points left of the identity line present sublinear summation, whereas any points right of the identity line are suggestive of supralinear summation. Figure 5B shows these plots for more than 20 branches for each neuron type. Across all neuron types, the majority of branches showed linear summation. RMSE values were less than 1.5 mV*s; thus, the measured response integrals were within 1.5 mV*s of the integral expected of linear summation. GM neurons (Figure 5Biv) exhibited particularly uniform linear summation across all branches evaluated and this is reflected in a small RMSE value of 0.32 mV*s. LP branches and PD branches show slightly higher RMSE values (greater than 1 mV*s), perhaps providing some evidence of variable shunting or amplifying mechanisms. Even so, RMSE values less than 2 mV*s across all cell types suggests relatively linear summation across most branches evaluated. Voltage summation arithmetic was similarly linear in the centrifugal direction (consult Supplements 1–4 to Figure 4). It should be noted that this evaluation does not consider the putative roles of TTX-sensitive sodium channels, which have been abolished with 10^−7^ M TTX in the bath.

**Figure 5.**
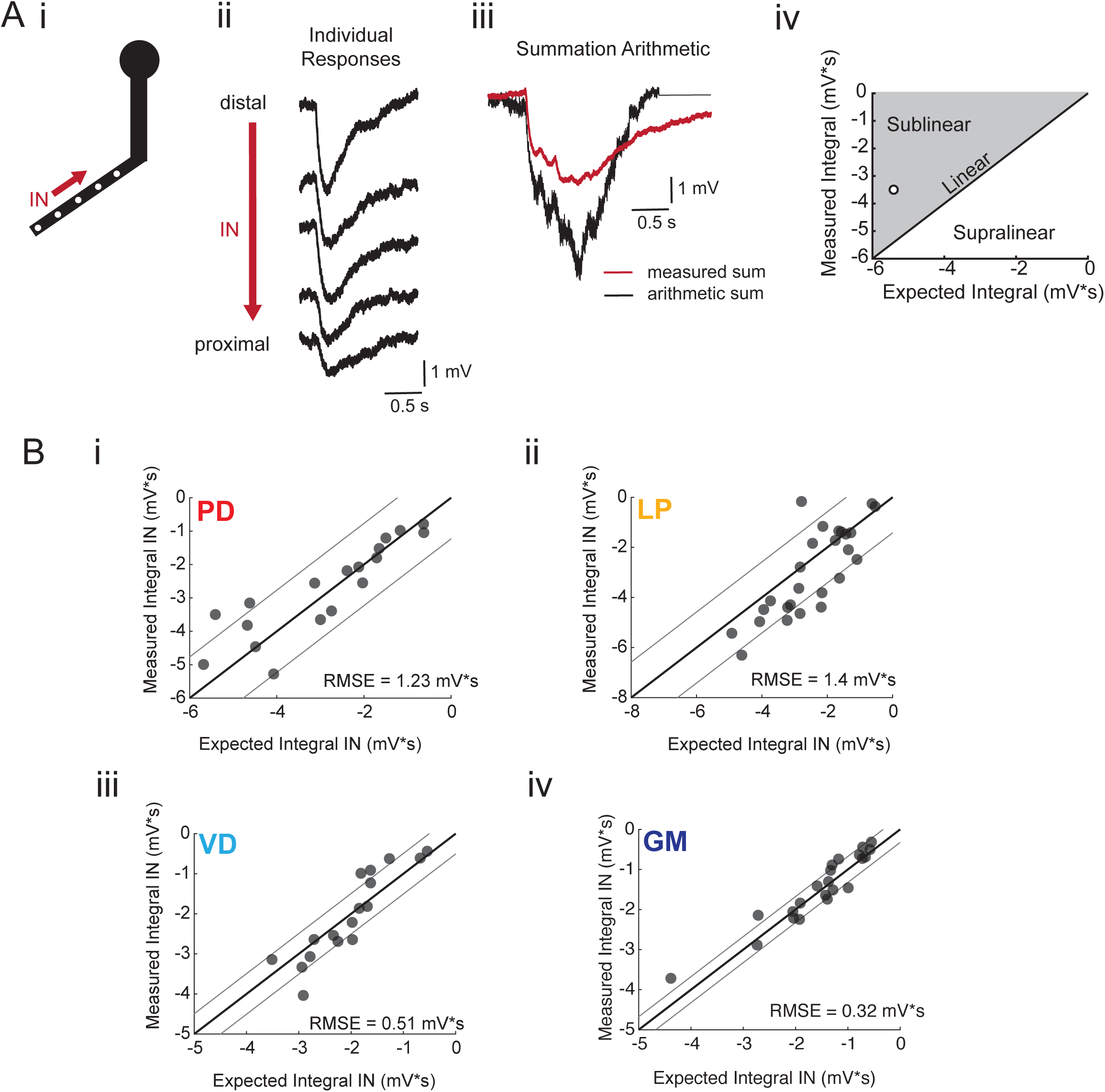
Arithmetic of voltage propagation in four neuron types. A. (i) 4–6 sites spaced 50–100 μm apart on the same secondary neurite were sequentially photoactivated at 5 Hz in the inward (IN) direction. (ii) Raw responses at individual sites from most distal (top) to most proximal (bottom) for one representative PD branch. (iii) Comparison of the inward voltage sum to the arithmetic sum for photo-activation of the individual sites (the traces shown in (ii) were summed with a 200-ms offset). (iv) The integrals (mV*s) of the measured responses were plotted against that of the expected arithmetic sum. This provides a graphical depiction of the linearity of voltage summation for each branch. Points to the left of the identity line suggest sublinear summation, points to the right of the identity line suggest supralinear summation, and points near or on the identity line suggest linear summation. The singular point depicted in (iv) depicts voltage summation for the responses shown in (iii). In this case, the measured voltage had a lesser integral than the arithmetic sum and therefore showed sublinear summation. B. (i-iv) Plots showing the measured integrals as a function of the expected integral for the arithmetic sum for the inward activation of many branches within the four neuron types: (i) 19 branches from 6 PD neurons, (ii) 27 branches from 9 LP neurons, (iii) 20 branches from 5 VD neurons, (iv) 22 branches from 5 neurons. These real data were fit to the identity line, and the root-mean-square error (RMSE) boundaries for this fit is plotted in gray lines.

## Discussion

Our proof-of-concept simulation demonstrated how voltage summation in single neurites is shaped by their biophysical and geometrical properties. This simulation recapitulated the canonical model wherein passive attenuation leads to directional selectivity and sublinear summation of sequentially evoked events on the same neurite (first presented by Rall (1964); shown in Figure 1A-C). Yet, by exploring a broader range of passive properties and neurite geometries, we showed that this dogmatic phenomenon may not hold true across the full range of geometrical features observed in diverse neuron types. Interestingly, we found that voltage summation in cable models with geometries reminiscent of those found in STG neurons, was directionally insensitive and linear for a broad range of passive properties. It is plausible that the neurite geometries observed in STG (Otopalik et al., 2017a) may contribute to the uniform and compact electrotonic structures observed in previous study (Otopalik et al., 2017b). To explore this possibility and its generalizability to all STG neuron types, we explored passive voltage signal propagation and summation in four STG neuron types using focal glutamate photo-uncaging.

### Interpreting electrotonus in STG neurons

In previous work, we showed that one neuron type, the Gastric Mill (GM) Neuron is electrotonically compact despite its expansive and complex morphology (Otopalik et al., 2017a, b). Here, we use similar experimental approaches to corroborate this result in the GM neuron, while also demonstrating that three other neuron types: PD, LP, and VD present similarly uniform and compact electrotonic structures, despite their different firing patterns and circuit functions. Small CV values for apparent E_rev_s measured in each neuron (Table 2) suggest uniformity of current flow across the neuronal structure. Furthermore, small CVs and mean E_rev_s between −60 and −80 mV within each neuron also suggest compactness of electrotonic structure, such that Erevs for responses evoked at the most distal sites do not undergo a substantial hyperpolarizing shift (a thorough discussion of this logic can be found in Otopalik et al., 2017b). Glutamate photo-uncaging across the neurite tree yielded somatic responses that varied in their magnitude. Thus, there is no evidence of distance-dependent scaling of receptors or other normalization mechanisms that have been observed in other neuron types (Magee and Cook, 2000; Andrásfalvy and Magee, 2001; Smith et al., 2003).

### Neurites with directional indifference

Our computational simulation revealed that the immense tapering of STG neurites may render them indifferent to direction of activation and yield linear summation. We tested this experimentally and observed no directional bias in any of the four neuron types; inhibitory potentials evoked in the inward and outward directions were similarly summed. Likewise, voltage summation was linear in most branches assessed. Thus, our experimental results align with the predictions of the model. Our simulation also suggested that the immense tapering of STG neurites allows for uniformity of voltage signal propagation across the neurite tree in the face of potential heterogeneity of passive biophysical properties. This result is interesting in light of the synaptic organization of the STG. Synaptic sites are sparsely distributed throughout the neuropil and pre- and post-synaptic sites are tightly apposed on the same neurites (King, 1976a, 1976b). Thus, sufficiently large synaptic potentials may originate anywhere on the neurite tree, propagate in any direction, and achieve consistent neuronal output (and therefore circuit function).

### A general solution for reliable pace-making physiology

The motor rhythm generated by the STG relies on slow oscillations and graded transmission (Eisen and Marder, 1982; Marder and Eisen, 1984; Maynard and Walton, 1975; Graubard et al., 1980; Manor et al., 1997, 1999). Here, we suggest that neurite geometry may aid STG neurons in generating consistent physiological output, in the face of variable magnitudes and distributions of intrinsic and synaptic properties across animals (Prinz et al., 2004; Schulz et al., 2006; Marder and Goaillard, 2006; Goaillard et al., 2009; Marder, 2011).

## Materials & Methods

### Animals and Dissections

Wild-caught adult male Jonah Crabs (*Cancer borealis*) were acquired and maintained by the Marine Resources Center at the Marine Biological Laboratories in Woods Hole, MA. Animals were maintained on a 12-hour dark/12-hour light cycle without food and in chilled natural seawater (10–13 deg C) in a 2000-liter tank at a density of no more than 30 crabs per tank. STG dissections were executed as in Otopalik et al., 2017b and as previously described (Gutierrez and Grashow, 2009) in saline solution (440 mM NaCl, 11 mM KCl, 26 mM MgCl_2_, 13 mM CaCl_2_, 11 mM Trizma base, 5 mM maleic acid, pH 7.4–7.6). The intact stomatogastric nervous system, including: two bilateral commissural ganglia, esophageal ganglion, and stomatogastric ganglion (STG), as well as the *lvn, mvn, dgn* were dissected from the animal’s foregut and pinned down in a Sylgard-coated petri dish (10 mL). The preparation was continuously superfused with chilled saline (11–13 degrees C) for the duration of the experiment using a bipolar temperature control system (Harvard Apparatus, CL-100).

### Electrophysiology and Dye-fills

All electrophysiology and dye-fill methods are consistent with those utilized in Otopalik et al., 2017b. The STG was desheathed for access to somata for intracellular recordings. These recordings were executed with glass micropipettes (20–30 MΩ) filled with internal solution: 10 mM MgCl_2_, 400 mM potassium gluconate, 10 mM HEPES buffer, 15 mM NaSO_4_, 20 mM NaCl (Hooper et al., 2015). Intracellular recordings signals were amplified with an Axoclamp 900A amplifier (Molecular Devices, as described in Otopalik et al. 2017b). For extracellular nerve recordings, Vaseline wells were built around the *lvn, mvn*, and *dgn* nerves and stainless-steel pin electrodes were used to monitor extracellular nerve activity (as indicated in Figure 1A). Extracellular nerve recordings were amplified using a Model 3500 extracellular amplifier (A-M Systems). All recordings were acquired with a Digidata 1550 (Molecular Devices) digitizer and visualized with pClamp data acquisition software (Axon Instruments, version 10.7). Neuron types were identified by matching concurrent intracellular spiking patterns with units on nerves known to contain their axons (as in Figure 1B) and verified with positive and negative current injections. After identification, a single neuron was filled with dilute alexa488 dye (2 mM Alexa Fluor 488-hyrazide sodium salt (ThermoFisher Scientific, catalog no. A-10436, dissolved in internal solution)) with negative current pulses (–4 nA, 500 ms at 0.5 Hz) for 15–25 min. Following the dye-fill, input resistance was measured at the soma in two-electrode current clamp (neurons with input resistances <5 MΩ were discarded). For two-electrode current clamp, the electrode containing dilute alexa488 was used for recording and amplified on a 0.1xHS headstage. The electrode used for cell identification was used for current injection and amplified with a 1xHS headstage. Input resistance was measured throughout the experiment and neurons with input resistances <5 MΩ were discarded. Reversal potentials for glutamate-evoked responses were determined by evoking responses at > eight membrane potentials between −100 and −40 mV. In some experiments, neurons were filled with 2% Lucifer Yellow CH dipotassium salt (LY; Sigma, catalog no. L0144; diluted in filtered water) for post-hoc imaging. LY was injected with a low-resistance (10–15 MΩ) glass micropipette for 20–50 min with negative current pulses (−6 to –8 nA, 500 ms at 0.5 Hz).

### Focal Glutamate Photo-uncaging

Focal glutamate photo-uncaging methodology was consistent with the methods used in Otopalik et al., 2017b, although different instrumentation was used. For photo-uncaging experiments, preparations were superfused with a re-circulating peristaltic pump to maintain a stable bath volume. 250 μM MNI-caged-L-glutamate (dissolved in saline; Tocris Bioscience, catalog no. 1490) was bath applied. 10^−7^ M teterodotoxin (TTX) was also superfused to minimize spike-driven synaptic activity. Alexa488-filled neurons were visualized and focal photo-uncaging was achieved with a Laser Applied Stimulation and Uncaging (LASU) system (Scientifica). In brief: this system was composed of an epifluorescence microscope (SliceScope, Scientifica) equipped with a 4x magnification air and 40x magnification water-immersion objective lenses (Olympus; PLN 4X and LUMPLFLN 40XW, respectively). A 780 IR-LED was used to visualize the stomatogastric ganglion and locate neurons of interest. A white fluorescence illumination system (CoolLED) and FITC/Alexa Fluor 488/Fluo3/Oregon Green filter set (Chroma) were used to excite and visualize fluorescent emission from neuronal dye-fills. Images were captured with a monochrome CCD camera (Scientifica, SciCam Pro; 1360x1025 array and 6.54 μm^2^ pixel size). Focal photo-activation of MNI-glutamate was achieved with a 405 nm laser (35 mW, spot size ≥ 1.5 μm with the 40x objective). The preparation platform and micromanipulators were mounted on a motorized movable base plate, allowing for smooth re-positioning (in the X-Y plane) of the objective over different neurites. For photo-activation at different sites within the field of view, the laser spot was re-positioned in quick sequence (5 Hz) using a set of X-Y galvanometers (Cambridge Technology, 6251H). Photo-uncaging sites within the field of view, laser pulse duration, and pulse rate were selected with the assistance of the LASU system software (Scientifica).

### Electrophysiology Analysis

Electrophysiological responses to focal glutamate photo-uncaging were visualized and analyzed offline, as previously described (Otopalik et al., 2017b), using a set of custom MATLAB (Mathworks, version 2017b) scripts that will be made available on the Marder Lab GitHub (https://github.com/marderlab).

### Post-hoc Imaging and Morphological Analysis

As in Otopalik et al., 2017b, each photo-uncaging site was re-located in fluorescence images of the alexa488 and/or Lucifer Yellow dye-fill acquired with the LASU microscope (at 40x and 20x magnification; described above). The distance between each site and the somatic recording site was measured utilizing a combination of 2-D and 3-D image stacks with the assistance of Simple Neurite Tracer on ImageJ/FIJI (Longair et al., 2011). Although most experiments were exclusively conducted using the LASU microscope, a subset of neurons were imaged at 40x magnification (Zeiss C-apochromat40x/1.2 W) on a VIVO microscope system equipped with a Yokogawa (CSU-X1, Japan) spinning disk confocal scan head mounted on a Zeiss Examiner microscope. Fluorescent dye-fills were visualized with standard GFP filters and images were captured with a Prime CMOS camera (Photometrics, 95B). Multiple image stacks in the z-dimension, spanning the STG, were stitched together with the assistance of the Stitching tool in ImageJ/FIJI (Preibisch et al., 2009).

### Passive cable models

The library of cable models utilized in Otopalik et al., 2017b was adapted to explore voltage summation in neurites with varying geometries (depicted in Figure 6) and passive properties: six specific axial resistances (R_a_): 10, 50, 100, 150, 200, 300 Ω*cm; and six specific membrane resistances (R_m_): 2 × 10^4^, 1.6 × 10^4^, 1 × 10^4^, 5 × 10^3^, 1 × 10^3^, 1 × 10^2^ Ω*cm^2^. All possible combinations of neurite geometries, membrane resistances, and axial resistances were assessed. All cables were 1000 μm in length and had a membrane capacitance of 1 μF*cm^−2^. Voltage summation was assessed by simulating our experimental procedure using the simulation platform NEURON (Hines and Carnevale, 2001). Voltage was recorded 100 μm from one end (d_0_) and inhibitory potentials (E_rev_ = −75 mV, *τ* = 70 ms, g_max_ = 10 nS) were evoked at five sites with increasing distance from the recording site (as depicted in Figure 1A). Sites were activated individually, and then at 5 Hz in the inward (toward the recording site) or outward (away from the recording site) directions. Using MATLAB (Mathworks, version 2017b), integrals were calculated for the inward and outward summed responses and the expected linear sum of the individual events in either direction. Directional bias was calculated as the inward integral minus the outward integral. Linearity was calculated as the expected arithmetic sum minus the recorded voltages sum for the inward direction.

**Figure 6.**
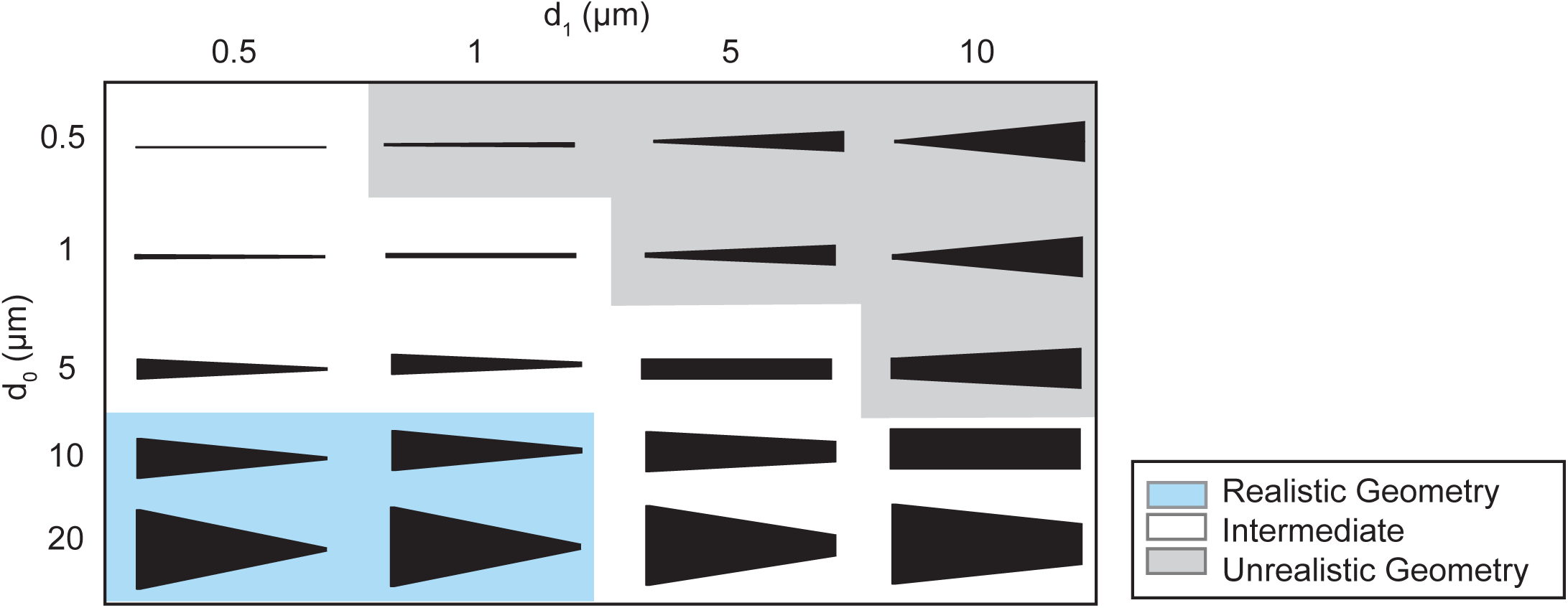
Schematic showing the range of cable geometries explored in the voltage summation simulation. d_0_ = diameter at recording site; d_1_ = diameter at distal end, 950 μm from the recording site. Shaded regions show qualitative relevance to the morphologies of actual secondary STG neurites (as measured in Otopalik et al., 2017a). These are indicated in the key.

## Acknowledgements

We thank Jennifer Bestman for assistance in spinning disk and confocal microscopy; the Marine Resources Center at the Marine Biological Laboratories for acquiring and maintaining animals; Louie Kerr at the Central Microscopy Facility; Dana Mock-Munoz de Luna for administrative support; Kamran Kodhakhah, Heather Rhodes, and the 2017 Grass Fellows for their support and feedback. This study was funded by the Grass Foundation and NINDS awards to EM and AO.

## Competing Interests

EM: Deputy editor, *eLife*. AO: No competing interests declared.

**Figure 1 - Supplement 1.**
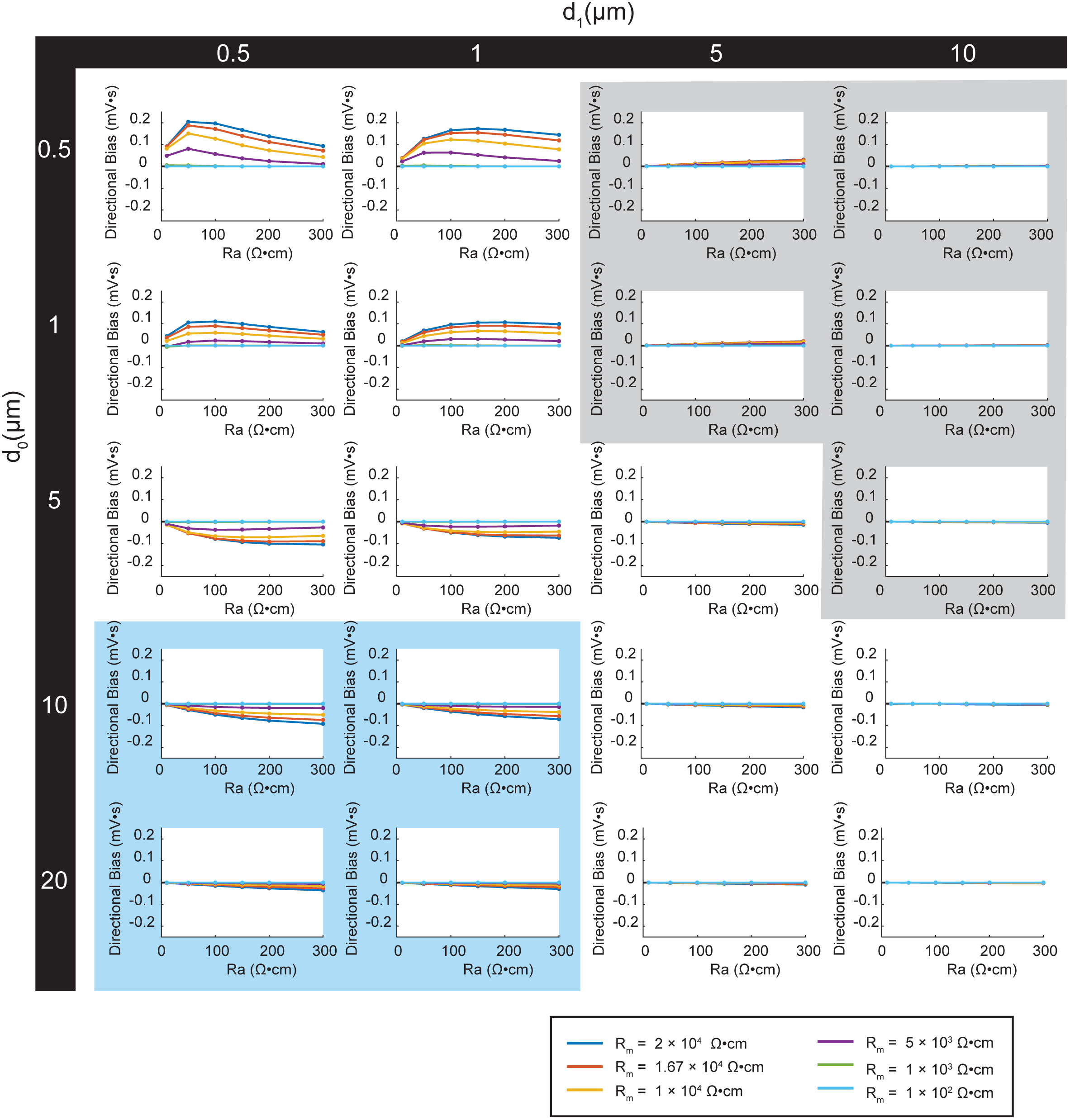
Simulating directional bias of voltage summation in neurites with diverse passive properties and geometries. Voltage summation experiments were simulated in NEURON as shown in Figure 6A. This array of plots shows directional bias, calculated as the integral of the inward voltage sum minus the outward voltage sum, as a function of specific axial resistance (R_a_ in Ω*cm) for a range of different specific membrane resistance (R_m_ in Ω*cm^2^) values (by color, shown in key). Positive data points suggest an inward bias, whereas negative values suggest an outward bias. Points close to y = 0 show no directional preference. Each column of plots corresponds to a different diameter at the distal end of the cable (d_1_), and each row of plots corresponds to a different diameter at the recording end of the cable (d_0_). These geometries are illustrated in Figure 6. The blue shaded region denotes neurite geometries with tapers reminiscent of those observed experimentally in secondary neurites of actual STG neurons (Otopalik et al., 2017a). The gray shaded region indicates neurite geometries that are unrealistic based on previous anatomical study of STG neurons.

**Figure 1 - Supplement 2.**
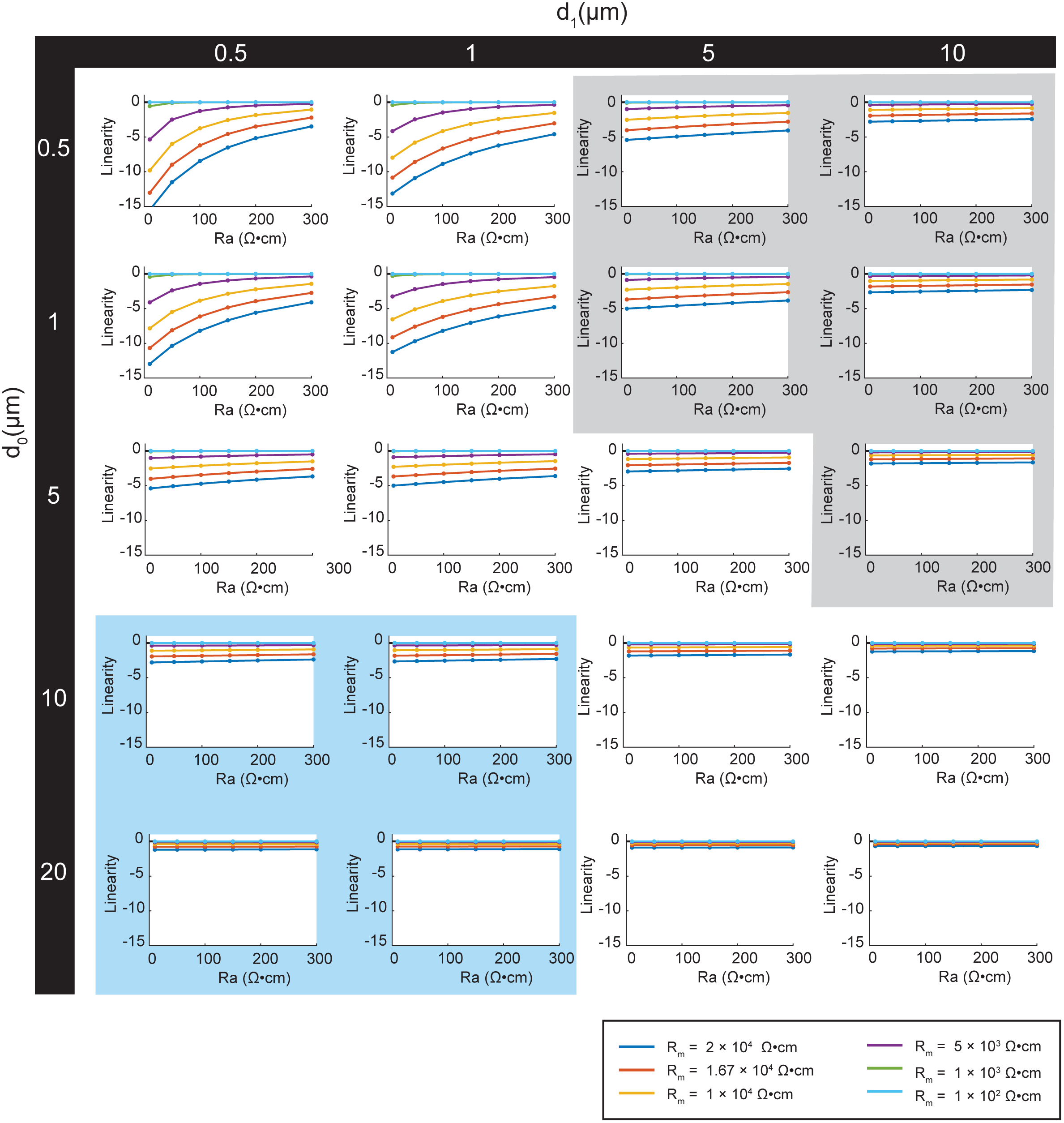
Simulating arithmetic of voltage summation in neurites with diverse passive properties and geometries. Voltage summation experiments were simulated in NEURON as shown in Figure 6A. This array of plots shows the linearity of voltage summation, calculated as the integral of the inward voltage sum minus the arithmetic sum of the individual events, as a function of specific axial resistance (R_a_ in Ω*cm) for a range of different specific membrane resistance (R_m_ in Ω*cm^2^) values (by color, shown in key). Positive data points suggest supralinear summation, whereas negative values suggest sublinear summation. Points close to y=0 indicate linear summation. Each column of plots corresponds to a different diameter at the distal end of the cable (d_1_), and each row of plots corresponds to a different diameter at the recording end of the cable (d_0_). These geometries are illustrated in Figure 7. The blue shaded region denotes neurite geometries with tapers reminiscent of those observed experimentally in secondary neurites of actual STG neurons (Otopalik et al., 2017a). The gray shaded region indicates neurite geometries that are unrealistic based on previous anatomical study.

**Figure 3 - Supplement 1.**
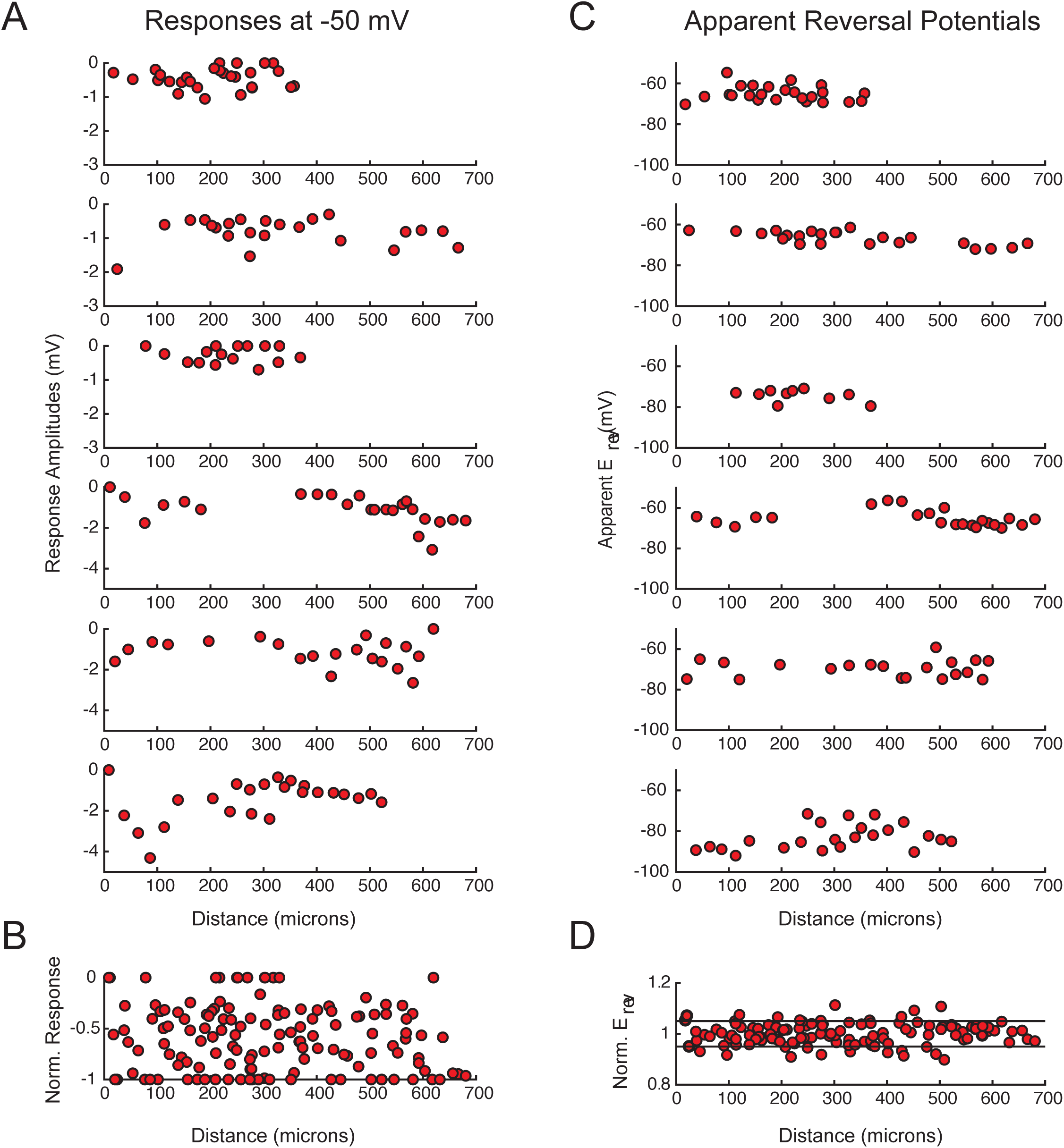
PD response magnitudes and apparent reversal potentials (Erevs) as a function of distance from the somatic recording site. A. Maximum response amplitudes were measured at −50 mV for individual sites. Each plot shows response amplitudes for many sites in one neuron. B. Apparent E_rev_s measured for each site. Each plot shows apparent E_rev_s for many sites in one neuron. B and D are the same normalized plots of response amplitude and apparent E_rev_s, respectively, shown in Figure 3.

**Figure 3 - Supplement 2.**
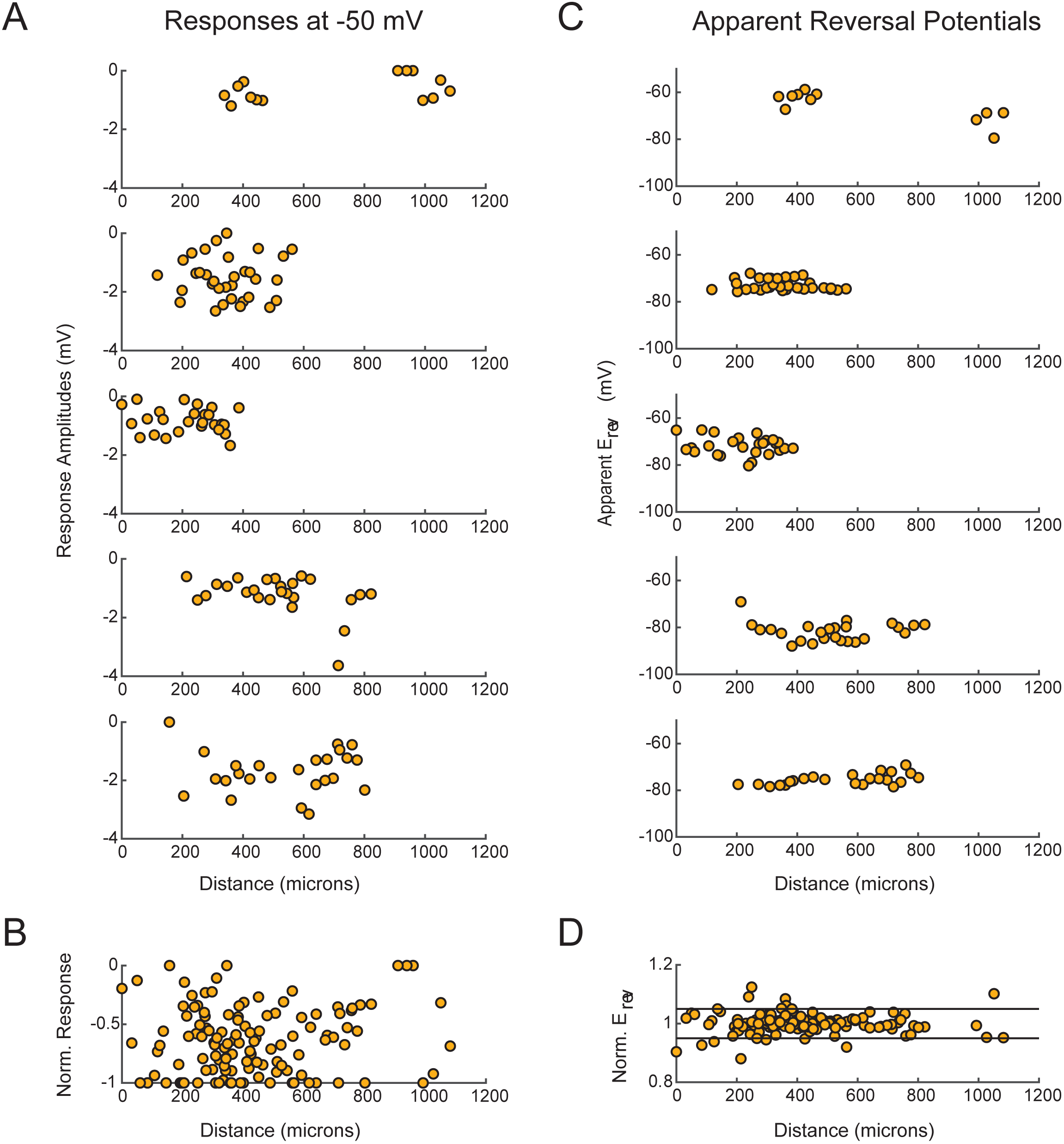
LP response magnitudes and apparent reversal potentials (E_rev_s) as a function of distance from the somatic recording site. A. Maximum response amplitudes were measured at −50 mV for individual sites. Each plot shows response amplitudes for many sites in one neuron. B. Apparent E_rev_s measured for each site. Each plot shows apparent E_rev_s for many sites in one neuron. B and D are the same normalized plots of response amplitude and apparent E_rev_s, respectively, shown in Figure 3.

**Figure 3 - Supplement 3.**
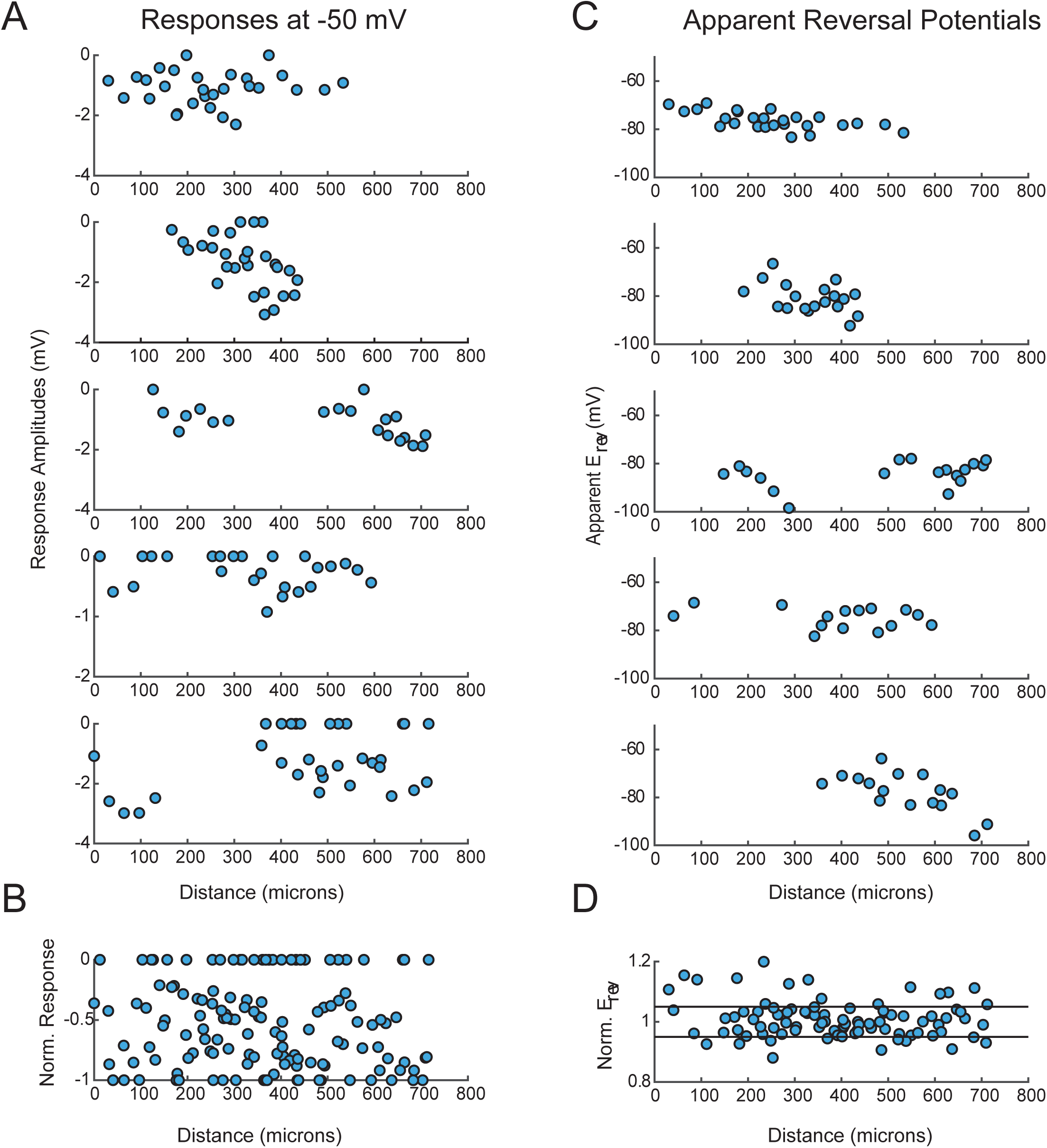
VD response magnitudes and apparent reversal potentials (E_rev_s) as a function of distance from the somatic recording site. A. Maximum response amplitudes were measured at −50 mV for individual sites. Each plot shows response amplitudes for many sites in one neuron. B. Apparent E_rev_s measured for each site. Each plot shows apparent E_rev_s for many sites in one neuron. B and D are the same normalized plots of response amplitude and apparent E_rev_s, respectively, shown in Figure 3.

**Figure 3 - Supplement 4.**
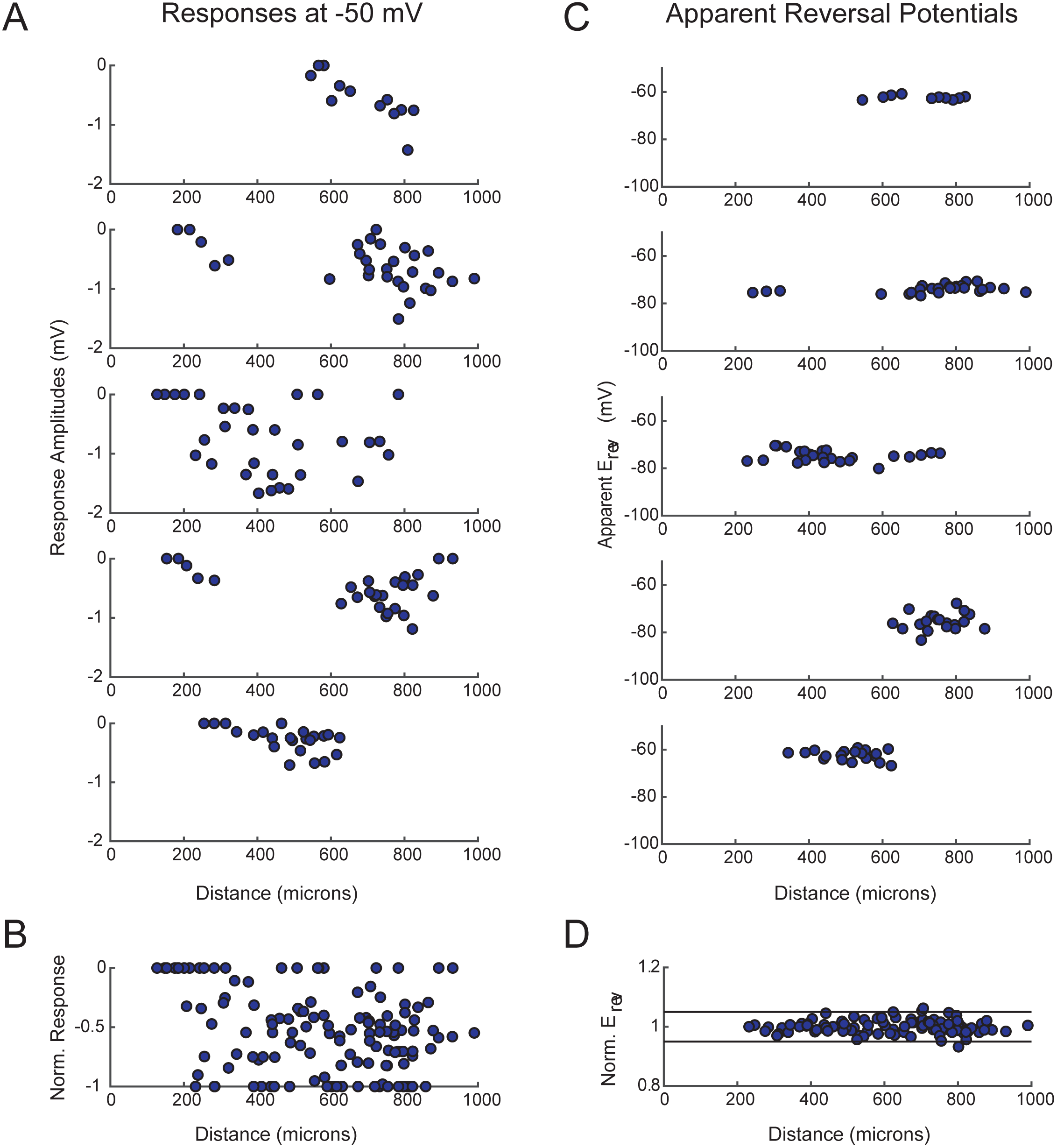
GM response magnitudes and apparent reversal potentials (E_rev_s) as a function of distance from the somatic recording site. A. Maximum response amplitudes were measured at −50 mV for individual sites. Each plot shows response amplitudes for many sites in one neuron. B. Apparent E_rev_s measured for each site. Each plot shows apparent E_rev_s for many sites in one neuron. B and D are the same normalized plots of response amplitude and apparent E_rev_s, respectively, shown in Figure 3.

**Figure 4 - Supplement 1.**
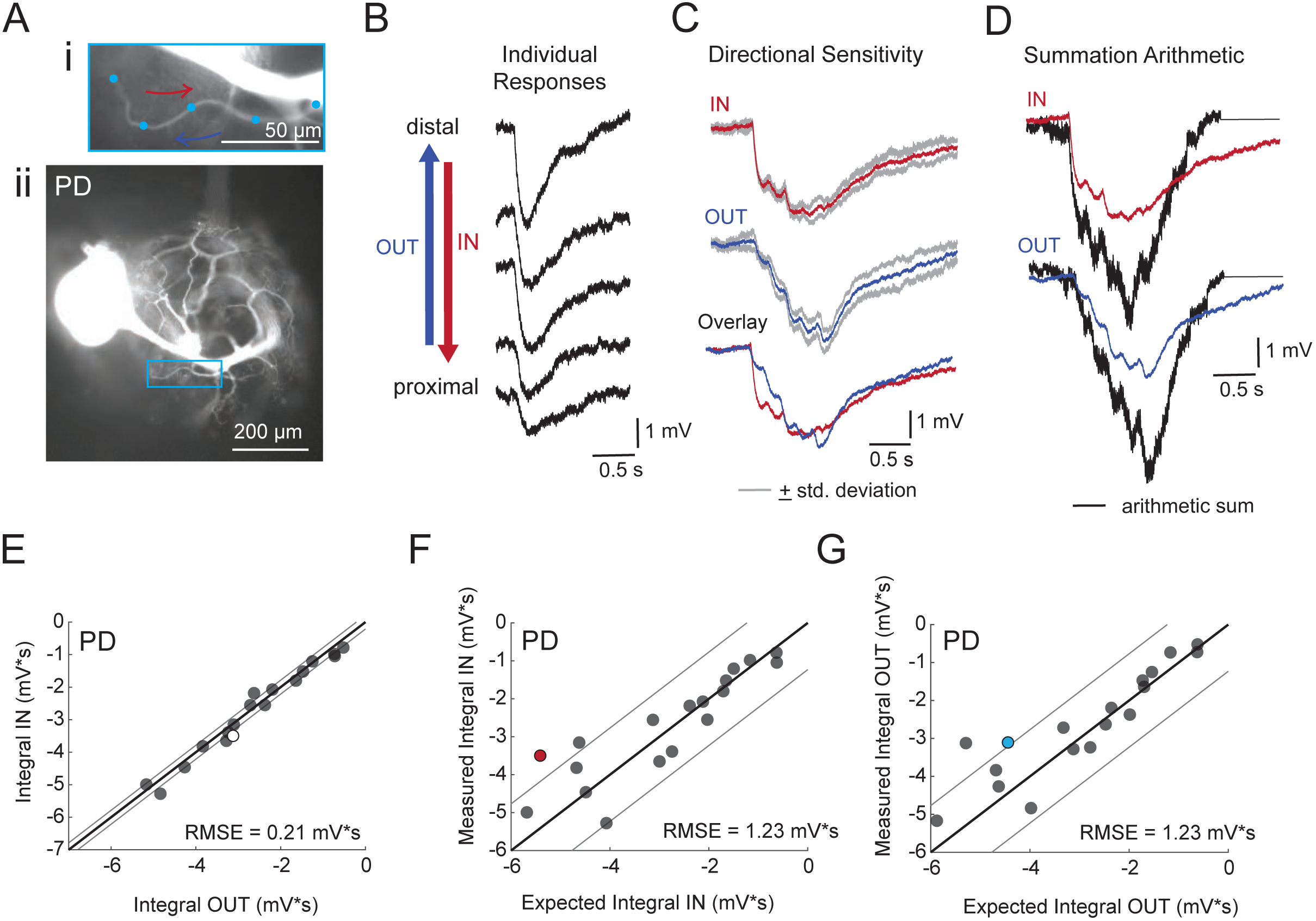
Directional sensitivity and arithmetic of voltage propagation in PD neurons. A. Representative images depicting one secondary neurite branch (i) at 40x magnification and (ii) in context of the entire dye-filled structure at 20x magnification. Arrows indicate inward (IN; red) and outward (OUT; blue) activation. B. Response evoked at −50 mV for individual sites shown in A from most distal site (top) to most proximal site (bottom). C. Comparison of mean summed voltage response for inward (red) and outward (blue) activation sequences at 5 Hz for the branch shown in A (mean calculated from 3 trials; the mean + standard deviation (gray) shows little variation across trials). D. Comparison of evoked responses (red, blue) to the arithmetic sum of the individual responses (summed with a 200-ms offset in either direction) for the branch in A. In this case, the measured responses are much less than the arithmetic sum expected for linear voltage summation. E. Plot of the measured inward integral (mV*s) as a function of the measured outward integral (mV*s). Points to the left of the identity line suggest an outward bias, points to the right of the identity line suggest an inward bias. Points near or on the identity suggest no directional preference. The white data point denotes the branch shown in part A. F-G: Plots of the measured integral (inward or outward, respectively, in mV*s) as a function of the integral expected from arithmetic summation of the responses at individual sites. Points to the left of the identity line suggest sublinear summation, points to the right of the identity line suggest supralinear summation. Points near or on the identity line suggest linear summation. Colored data points denote the data from the branch shown in A. In E-G: Show data for 19 branches from 6 PD neurons; RMSE boundaries are plotted as gray lines and show the goodness of fit to the identity line.

**Figure 4 - Supplement 2.**
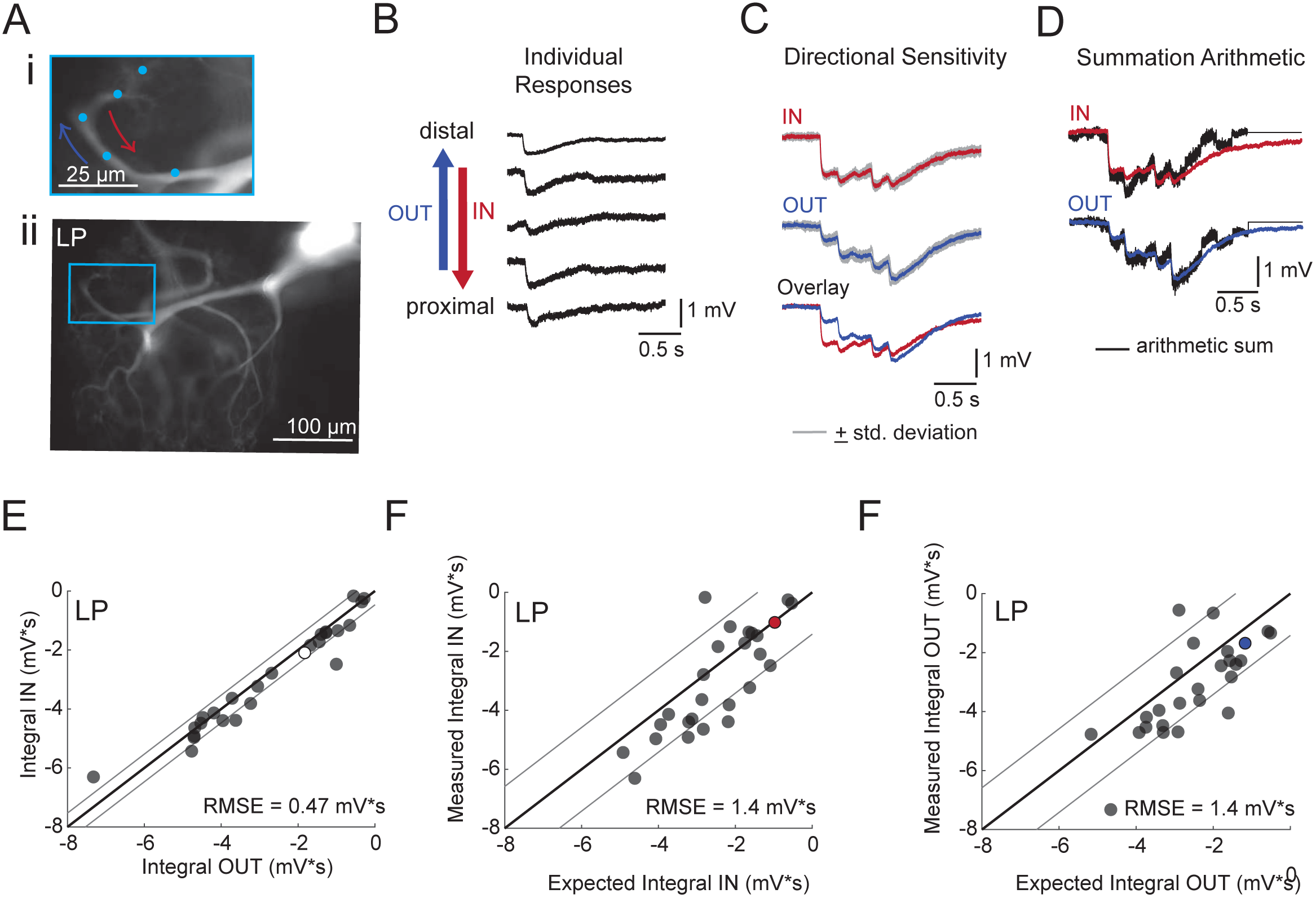
Directional sensitivity and arithmetic of voltage propagation in LP neurons. A. Representative images depicting one secondary neurite branch (i) at 40x magnification and (ii) in context of the entire dye-filled structure at 20x magnification. Arrows indicate inward (IN; red) and outward (OUT; blue) activation. B. Response evoked at −50 mV for individual sites shown in A from most distal site (top) to most proximal site (bottom). C. Comparison of mean summed voltage response for inward (red) and outward (blue) activation sequences at 5 Hz for the branch shown in A (mean calculated from 3 trials; the mean + standard deviation (gray) shows little variation across trials). D. Comparison of evoked responses (red, blue) to the arithmetic sum of the individual responses (summed with a 200-ms offset in either direction) for the branch in A. In this case, the measured responses are similar to the arithmetic sum. E. Plot of the measured inward integral (mV*s) as a function of the measured outward integral (mV*s). Points to the left of the identity line suggest an outward bias, points to the right of the identity line suggest an inward bias. Points near or on the identity suggest no directional preference. The white data point denotes the branch shown in part A. F-G: Plots of the measured integral (inward or outward, respectively, in mV*s) as a function of the integral expected from arithmetic summation of the responses at individual sites. Points to the left of the identity line suggest sublinear summation, points to the right of the identity line suggest supralinear summation. Points near or on the identity line suggest linear summation. Colored data points denote the data from the branch shown in A. In E-G: Show data for 26 branches from 9 LP neurons; RMSE boundaries are plotted as gray lines and show the goodness of fit to the identity line.

**Figure 4 - Supplement 3.**
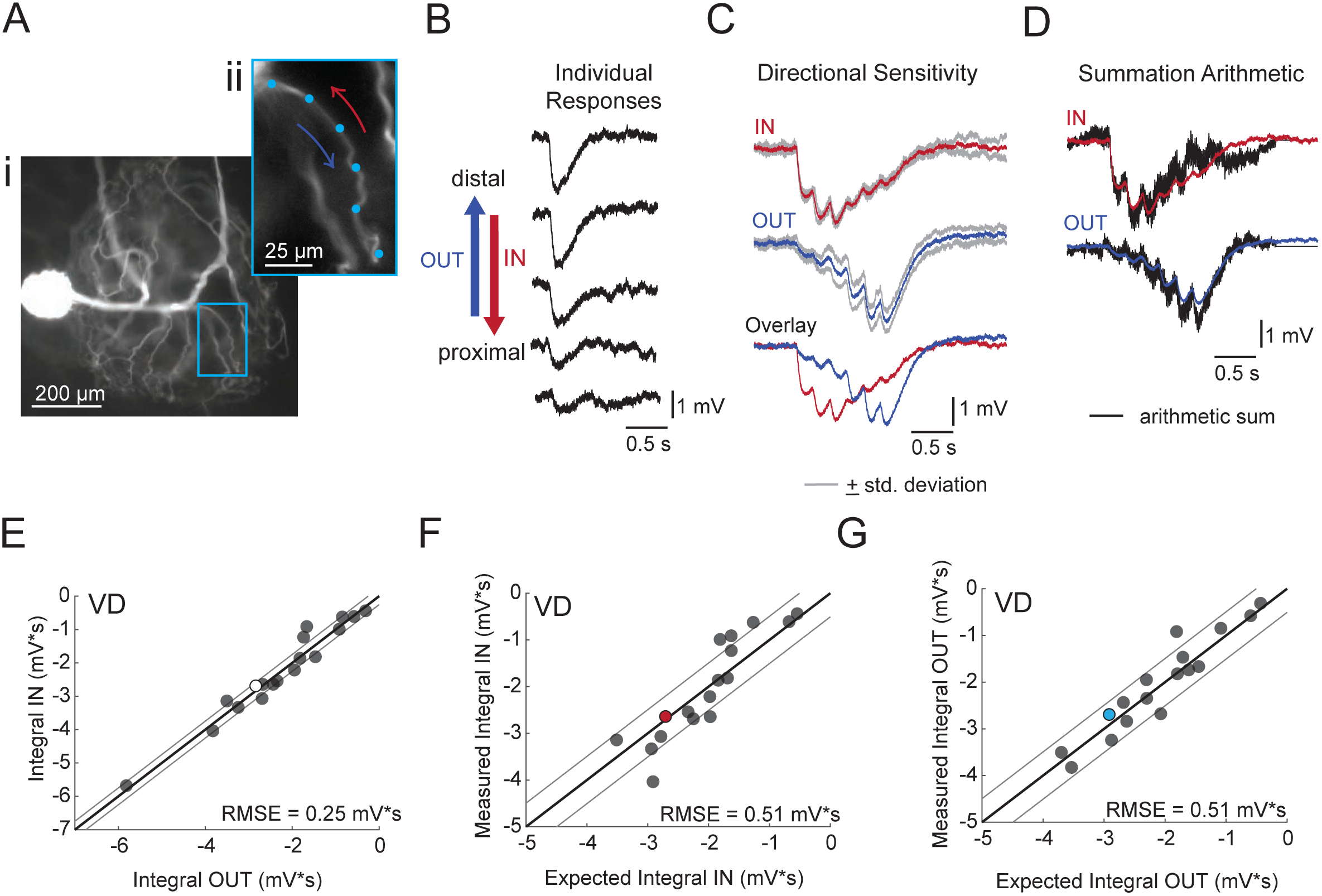
Directional sensitivity and arithmetic of voltage propagation in VD neurons. A. Representative images depicting one secondary neurite branch (i) at 40x magnification and (ii) in context of the entire dye-filled structure at 20x magnification. Arrows indicate inward (IN; red) and outward (OUT; blue) activation. B. Response evoked at −50 mV for individual sites shown in A from most distal site (top) to most proximal site (bottom). C. Comparison of mean summed voltage response for inward (red) and outward (blue) activation sequences at 5 Hz for the branch shown in A (mean calculated from 3 trials; the mean + standard deviation (gray) shows little variation across trials). D. Comparison of evoked responses (red, blue) to the arithmetic sum of the individual responses (summed with a 200-ms offset in either direction) for the branch in A. In this case, the measured responses are similar to the arithmetic sums. E. Plot of the measured inward integral (mV*s) as a function of the measured outward integral (mV*s). Points to the left of the identity line suggest an outward bias, points to the right of the identity line suggest an inward bias. Points near or on the identity suggest no directional preference. The white data point denotes the branch shown in part A. F-G: Plots of the measured integral (inward or outward, respectively, in mV*s) as a function of the integral expected from arithmetic summation of the responses at individual sites. Points to the left of the identity line suggest sublinear summation, points to the right of the identity line suggest supralinear summation. Points near or on the identity line suggest linear summation. Colored data points denote the data from the branch shown in A. In E-G: Show data for 18 branches from 5 VD neurons; RMSE boundaries are plotted as gray lines and show the goodness of fit to the identity line.

**Figure 4 - Supplement 4.**
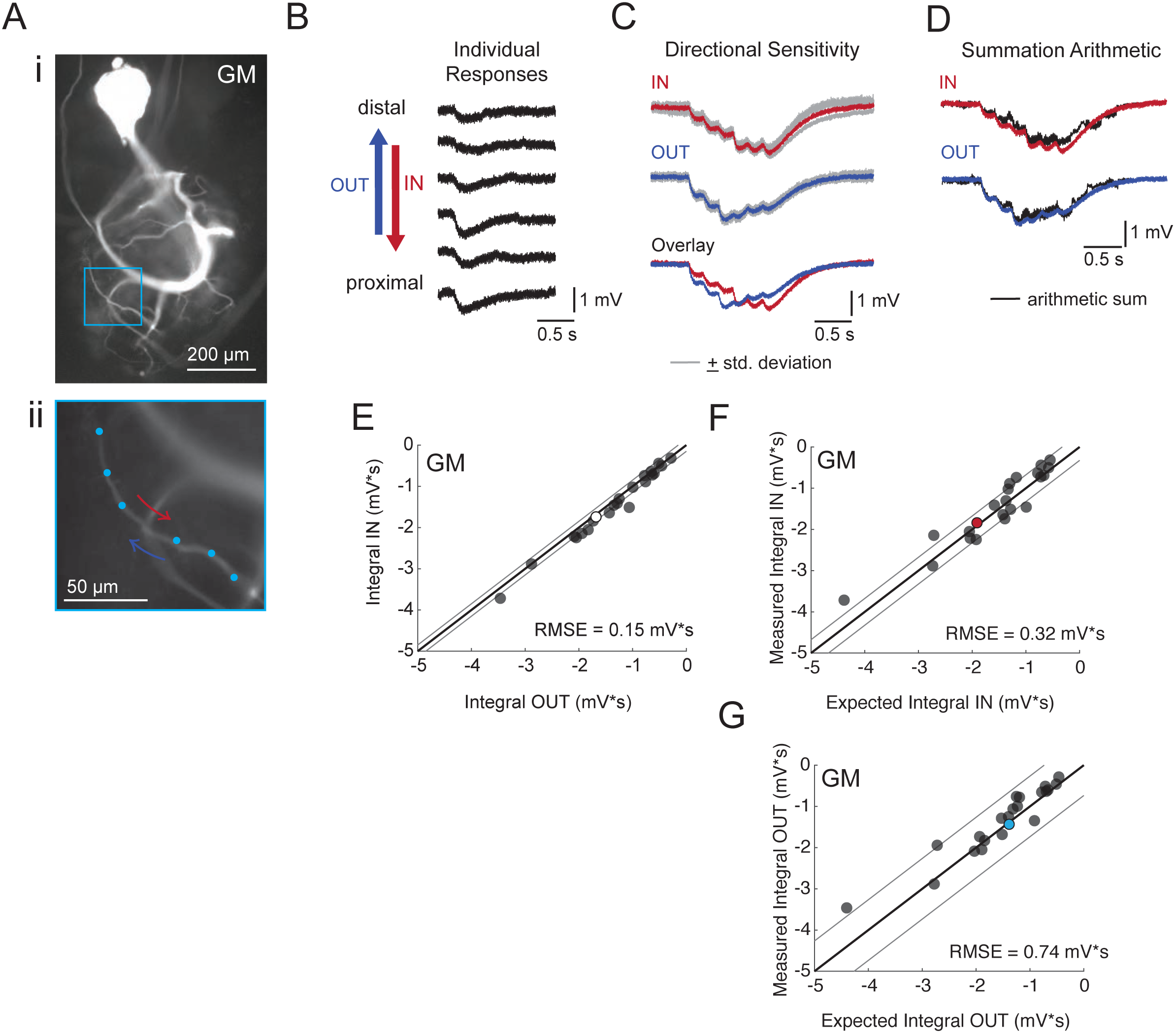
Directional sensitivity and arithmetic of voltage propagation in GM neurons. A. Representative images depicting one secondary neurite branch (i) at 40x magnification and (ii) in context of the entire dye-filled structure at 20x magnification. Arrows indicate inward (IN; red) and outward (OUT; blue) activation. B. Response evoked at −50 mV for individual sites shown in A from most distal site (top) to most proximal site (bottom). C. Comparison of mean summed voltage response for inward (red) and outward (blue) activation sequences at 5 Hz for the branch shown in A (mean calculated from 3 trials; the mean + standard deviation (gray) shows little variation across trials). D. Comparison of evoked responses (red, blue) to the arithmetic sum of the individual responses (summed with a 200-ms offset in either direction) for the branch in A. In this case, the measured responses are similar to the arithmetic sums. E. Plot of the measured inward integral (mV*s) as a function of the measured outward integral (mV*s). Points to the left of the identity line suggest an outward bias, points to the right of the identity line suggest an inward bias. Points near or on the identity suggest no directional preference. The white data point denotes the branch shown in part A. F-G: Plots of the measured integral (inward or outward, respectively, in mV*s) as a function of the integral expected from arithmetic summation of the responses at individual sites. Points to the left of the identity line suggest sublinear summation, points to the right of the identity line suggest supralinear summation. Points near or on the identity line suggest linear summation. Colored data points denote the data from the branch shown in A. In E-G: Show data for 22 branches from 5 VD neurons; RMSE boundaries are plotted as gray lines and show the goodness of fit to the identity line.

